# Confined keratocytes mimic *in vivo* migration and reveal volume-speed relationship

**DOI:** 10.1101/2022.07.19.500705

**Authors:** Ellen C. Labuz, Matthew J. Footer, Julie A. Theriot

## Abstract

Fish basal epidermal cells, known as keratocytes, are well-suited for cell migration studies. *In vitro*, isolated keratocytes adopt a stereotyped shape with a large fan-shaped lamellipodium and a nearly spherical cell body. However, in their native *in vivo* environment, these cells adopt a significantly different shape during their rapid migration towards wounds. Within the epidermis, keratocytes experience 2D confinement between the outer epidermal cell layer and the basement membrane; these two deformable surfaces constrain keratocyte cell bodies to be flatter *in vivo* than in isolation. *In vivo* keratocytes also exhibit a relative elongation of the front-to-back axis and substantially more lamellipodial ruffling, as compared to isolated cells. We have explored the effects of 2D confinement, separated from other *in vivo* environmental cues, by overlaying isolated cells with an agarose hydrogel with occasional spacers, or with a ceiling made of PDMS elastomer. Under these conditions, isolated keratocytes more closely resemble the *in vivo* migratory shape phenotype, displaying a flatter apical-basal axis and a longer front-to-back axis than unconfined keratocytes. We propose that 2D confinement contributes to multiple dimensions of *in vivo* keratocyte shape determination. Further analysis demonstrates that confinement causes a synchronous 20% decrease in both cell speed and volume. Interestingly, we were able to replicate the 20% decrease in speed using a sorbitol hypertonic shock to shrink the cell volume, which did not affect other aspects of cell shape. Collectively, our results suggest that environmentally imposed changes in cell volume may influence cell migration speed, potentially by perturbing physical properties of the cytoplasm.

## Introduction

After injury to an animal’s skin, the wound-healing response restores tissue architecture and protects against entry of pathogens. Multiple cell types orchestrate an array of responses depending on the nature of the wound, including immune defense, cell proliferation, and production of extracellular matrix (Martin, 1997; Singer & Clark, 1999). The restoration of epithelium over the wound, or re-epithelialization, is critical for barrier function across animal taxa. Mammalian re-epithelialization requires that basal keratinocytes proliferate near the wound margin and migrate under the desiccated scab using actin-based lamellipodia (Singer & Clark, 1999). The multi-layered adult zebrafish epidermis heals using basal cell crawling and intercalation events (Richardson et al., 2016). In embryonic epithelia, from insects to mammals, rapid wound closure is driven by actomyosin “purse-string” contraction at the wound margin, often in combination with lamellipodial motility (Abreu-Blanco et al., 2012; McCluskey & Martin, 1995; Radice, 1980).

*In vivo* work has suggested that re-epithelialization occurs in a dynamic mechanical environment, yet many studies of the biophysical mechanisms driving wound closure have occurred outside of the relevant tissue context. Depending on the species of animal and the type of injury, the initial wound and subsequent healing process can lead to changes in tissue tension, extracellular matrix composition, and cell crowding (Evans et al., 2013; Franco et al., 2019; Singer & Clark, 1999). In contrast, studies of cultured cells on plastic or glass surfaces provide far simpler environments. Such studies illustrate how molecular interactions determine cell-scale forces and ultimately tissue-scale stresses and motion (Mogilner et al., 2020; Trepat & Fredberg, 2011). *In vitro* culture systems, from simple to complex, can also reveal cell-intrinsic responses to environmental complexities by allowing independent control over many external variables. Given the recent proliferation of methods for simulating physiological microenvironments, we were particularly interested in identifying physical properties of wounds and the wound environment that promote or limit efficient migration of epidermal cells.

Zebrafish provide an excellent means to study the same cell type migrating during wound healing both *in vivo* and in culture. Re-epithelialization has been particularly well studied at the larval stage, because of the organism’s optical clarity, extremely fast healing response (<30 minutes), and simple two-layered epidermal architecture (Arora et al., 2020; Kennard & Theriot, 2020; Rasmussen et al., 2015; Sonawane et al., 2005). The outer periderm cell layer maintains electrochemical gradients via tight junctions and forms an actomyosin purse-string that contracts the wound margin, while the inner basal cells serve as epidermal stem cells and crawl rapidly towards the wound on a basement membrane using actin-rich lamellipodia (Gault et al., 2014; Lee et al., 2014; Sonawane et al., 2005). Physical cues for basal cell migration include osmotic shock and electric fields. These develop locally when skin wounds disrupt transepithelial gradients of osmotic pressure and ionic electrical potential (Gault et al., 2014; Kennard & Theriot, 2020).

In addition to contributing to the wound response *in vivo*, zebrafish skin cells are also well-studied in culture. Isolated fish basal epidermal cells, known as keratocytes, move rapidly and persistently while maintaining a remarkably constant fan-like shape (Goodrich, 1924). This simple cell-scale geometry has facilitated many quantitative studies on cytoskeletal mechanisms of motility and shape determination (Mogilner et al., 2020). Polymerization of a dense actin network at the leading edge smoothly advances the membrane (Garner & Theriot, 2022; Keren et al., 2008; Theriot & Mitchison, 1991), while contraction of non-muscle myosin II at the rear powers actin retrograde flow, network compaction, and actin disassembly (Svitkina et al., 1997; Wilson et al., 2010). To transmit these forces to the substrate, adhesions are located throughout the lamellipodium and are particularly prominent at the cell’s rear corners (Fournier et al., 2010; Oliver et al., 1999). Physical perturbations, such as changes to substrate adhesiveness and membrane area, directly affect keratocyte shape and speed (Barnhart et al., 2011; Lieber et al., 2013).

Together, *in vivo* and isolated keratocyte studies can be used in complementary ways to examine how the physical environment of a wound affects migration. For example, *in vivo* keratocytes are confined to a 2D plane, sandwiched between a basement membrane and the periderm. While the forces exerted by these tissue layers are not known, they are notably absent for isolated keratocytes under standard culture conditions where individual cells move freely on a rigid substrate such as glass or polystyrene. Therefore, isolated keratocytes can be examined with and without experimentally applied confinement in order to simulate, and thereby better understand, *in vivo* skin conditions. Interestingly, a number of confinement methods that restrict cells to a thin 2D plane have recently been deployed for a variety of other cell types in culture, revealing surprising changes in behavior, as compared to unconfined conditions. These changes include the formation of stable bleb-based protrusions, dorsal actin patches, and seemingly unproductive rear blebs and dorsal adhesions (Gaertner et al., 2022; George et al., 2018; Ramalingam et al., 2015; Rape & Kumar, 2014; Srivastava et al., 2020). 2D confinement has also been shown to induce dramatically faster migration, particularly in transformed and poorly-adherent cells (Bergert et al., 2012; Liu et al., 2015; Logue et al., 2015; Ruprecht et al., 2015, 2015; Toyjanova et al., 2015; Tozluoğlu et al., 2013). However, less is known about the effect of 2D confinement on adherent, well-polarized cell types, especially cells that are already adapted to a flat, thin environment, such as the fish keratocyte.

Comparing *in vivo* and isolated keratocyte cell shapes at high spatiotemporal resolution during rapid migration, we find that keratocytes adopt a taller, rounder cell body in the absence of the skin’s physical constraints. Simply adding 2D confinement to isolated cells, via multiple methods, achieved good mimicry of the overall cell shapes observed *in vivo*. Furthermore, our well-controlled confinement system allowed a detailed quantification of keratocyte shape, size, and speed during a sudden change in environment, revealing an unexpected relationship between volume loss and speed decrease.

## Results

### Keratocytes experience 2D physical constraints in the zebrafish epidermis

In order to understand the role of the physical environment in keratocyte migration, we compared cells from the same lineage migrating in two settings: 1) the caudal fin epidermis after wounding by laceration (Kennard et al., 2021), and 2) primary culture after epidermal dissociation (Lou et al., 2015). We conducted time-lapse confocal microscopy on *in vivo* cells using wounded larvae expressing LifeAct-mNeonGreen in a mosaic fashion in approximately 5% of cells (*Methods*). Laceration was performed in order to induce basal cell migration at 3 days post fertilization (dpf), when larvae have naturally emerged from their chorion and are well-sized for manual wounding. In parallel, we also examined migration of isolated cells from embryos at a similar developmental stage using primary culture from a stable LifeAct-EGFP fish line. Keratocytes were extracted from 2 dpf larvae to obtain the optimal number of polarized cells (Lou et al., 2015).

These movies confirmed that keratocytes in both conditions migrate rapidly with an actin-rich lamellipodium (**Figure 1**, arrowheads). *In vivo,* cells migrate collectively towards the wound for a relatively short duration, previously measured to be less than 15 minutes in our standard tissue laceration (Kennard & Theriot, 2020). In contrast, isolated cells migrate persistently for up to several hours without any directional cue, as previously observed (Goodrich, 1924). Using apical-basal cross-sections, we observed that *in vivo* cells have a uniform height of between 3-4 µm (**Figure 1A, Video S1**). In contrast, isolated cells have a spherical cell body situated behind the thin lamellipodium, where the spherical body can be very tall (∼8 µm) while the lamellipodium is typically more uniform and much flatter than its counterpart in vivo (<0.5 µm) (**Figure 1B, Video S2**). We also observed that cells migrating *in vivo* are elongated parallel to the direction of migration and exhibit lamellipodial ruffling, but isolated cells are wide with a smooth fan-shaped lamellipodium and their longest axis perpendicular to the direction of migration (**Figure 1C**). Since keratocyte cell bodies adopt a taller, rounder shape without the influence of the epidermal environment, we concluded that the overlying cell layer and the basement membrane place a geometric constraint on basal cells, confining them two-dimensionally. We hypothesized that this physical confinement, rather than specific cell-cell interactions or soluble factors, could be largely responsible for the shape differences between *in vivo* and isolated basal cells.

**Figure 1.**
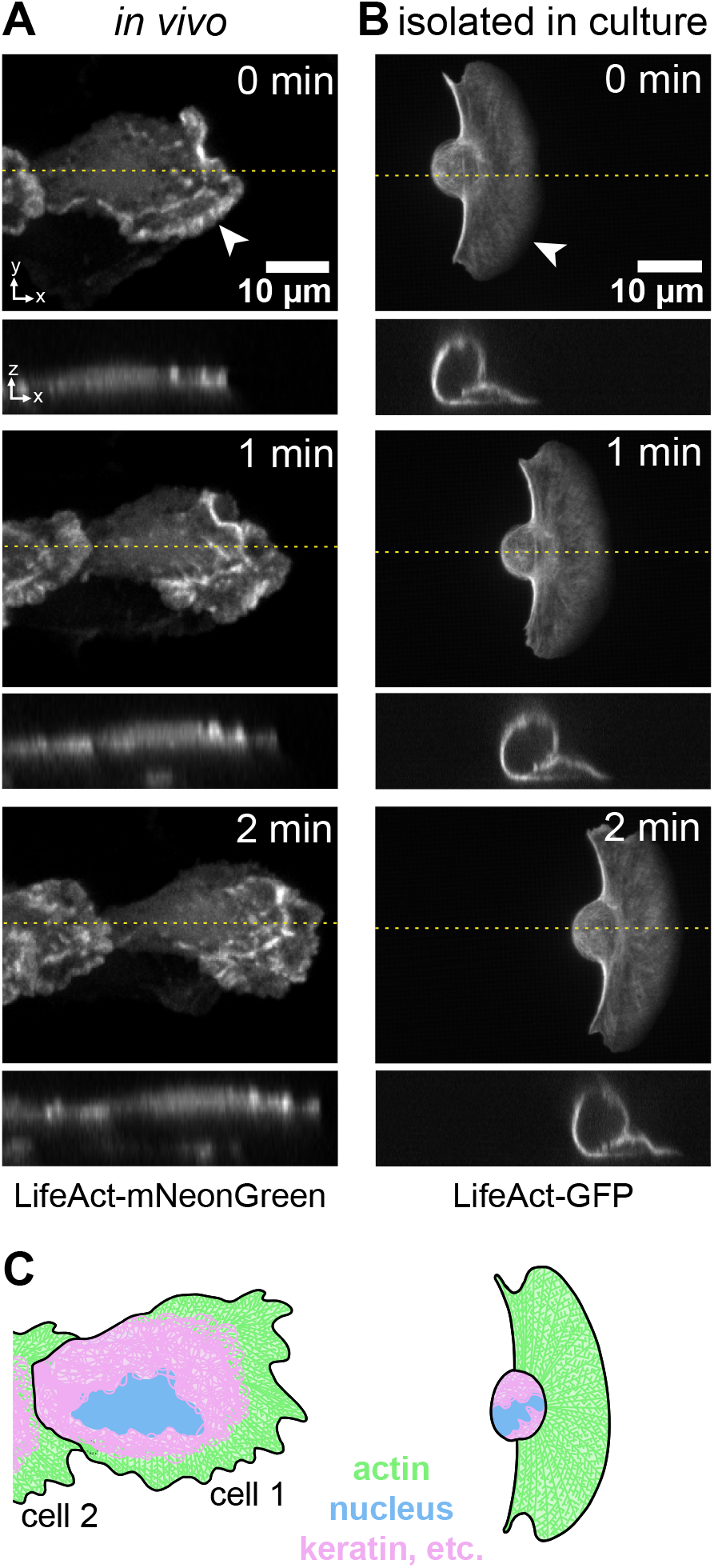
Keratocytes experience 2D physical constraints in the zebrafish epidermis. **(A)** Individual cell over time from a wounded 3 days post fertilization (dpf) larva expressing LifeAct-mNeonGreen mosaically in keratocytes (*TgBAC(ΔNp63:Gal4)* larvae injected with *UAS:LifeAct-mNeonGreen-P2A-mRuby3-CAAX* plasmid at the 1- or 2-cell stage). Laceration wound was to the right approximately 3 minutes earlier. **(B)** Isolated cell over time extracted from 2 dpf larvae expressing LifeAct-EGFP in keratocytes (*TgBAC(ΔNp63:Gal4)*; *Tg(UAS:LifeAct-EGFP))*. Extraction was approximately 2.5 hours earlier onto a collagen-coated coverslip. Images are gamma-corrected to better visualize the entire cell. Arrowheads: actin-rich lamellipodia. For **(A,B)**, XY-plane images are maximum-intensity Z-projections from spinning-disk confocal stacks. XZ-planes are side views taken at the position indicated by the dashed yellow line.

### Isolated keratocytes adopt elongated, narrow shapes under two methods of 2D confinement

We predicted that the introduction of artificial 2D confinement could induce an overall shape change in isolated cells, rendering them more similar in their shape and organization to cells observed *in vivo*. We sought to directly compare isolated keratocytes in confined and unconfined environments using multiple confinement strategies. First, we confined isolated keratocytes beneath an agarose gel, along with rigid polystyrene beads. Each bead held the agarose gel above the substrate in its local area like a tent-pole. This geometry made it possible for motile cells to move reversibly from areas of strict confinement to areas without confinement over short distances in a single field of view (**Figure 2A**). These non-confined areas helped us to control for gel effects other than physical contact. Using phase contrast imaging, we observed that cells far from beads, presumably confined by the agarose, displayed unusual shapes for isolated keratocytes. The confined cell shapes featured an ovoid cell body, elongated parallel to the direction of migration and positioned behind the lamellipodium (**Figure 2B**, top panel). For the cell shown in Figure 2B, we recorded its migration as it transitioned from a confined region to a region near an agarose-elevating bead, where the cell displayed a wider phase halo that encircled the entire cell body, suggesting increased height as expected (**Figure 2B**, yellow arrowhead). As the cell became less confined, it simultaneously adopted a more circular cell body that rested on top of a wider lamellipodium, similar to keratocytes in unconfined samples (**Figure 2B,C, Video S3**). These observations suggest that 2D confinement is directly connected to an elongated, narrow cell shape, with confined cells elongating parallel to their axis of migration and unconfined cells generating wider lamellipodia such that the cell’s major axis becomes perpendicular to its direction of migration. These observations further indicated that the shape changes between these two states were rapidly reversible over 1-2 minutes and depend only on the cell’s immediate environment and not on its recent history.

**Figure 2.**
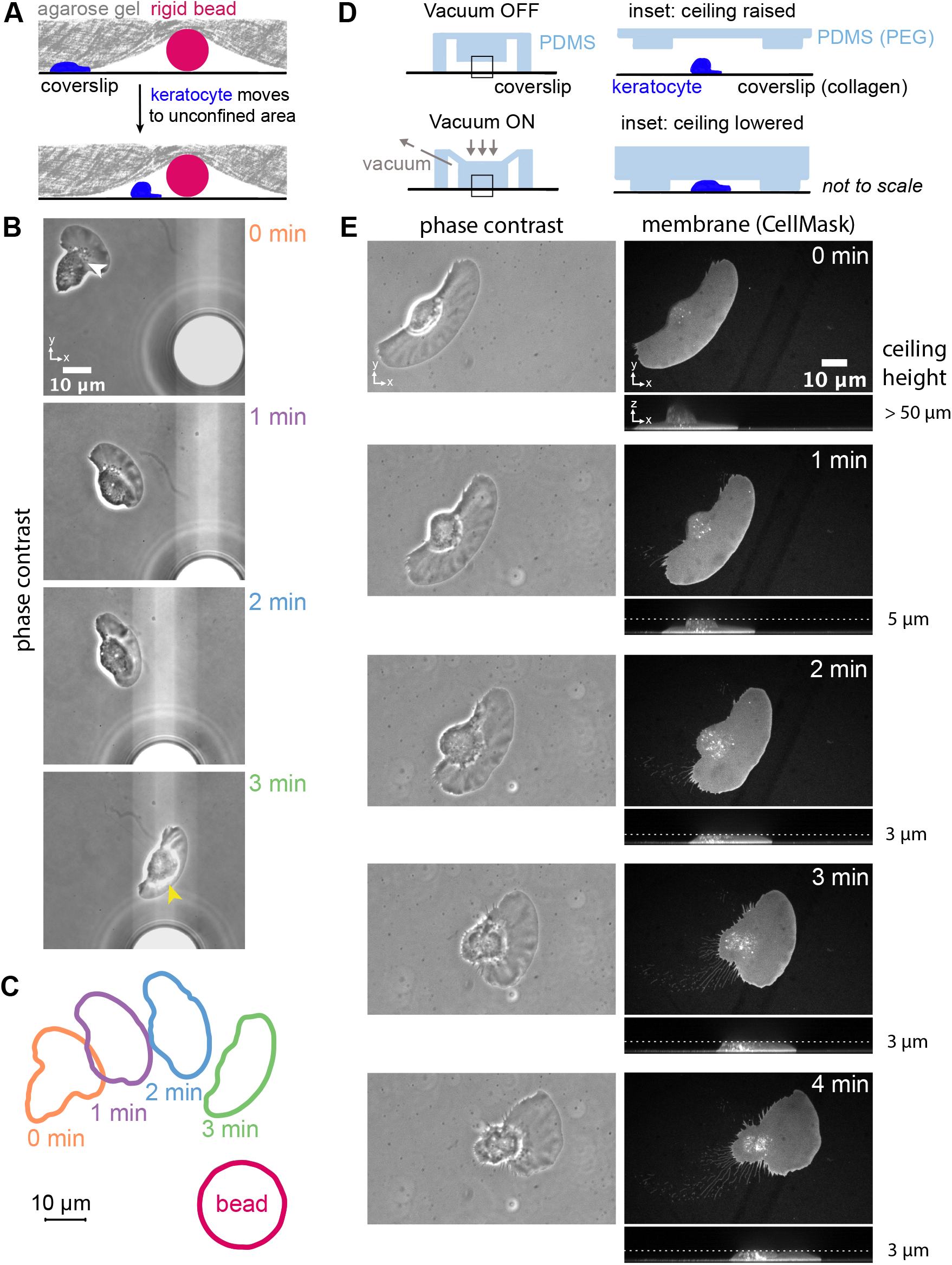
Isolated keratocytes adopt elongated, narrow shapes under two methods of 2D confinement. **A** Schematic of a rigid polystyrene bead propping up agarose like a tent-pole to create a local area without confinement. **(B)** Keratocyte that moved from a gel-confined area (1% agarose) to an unconfined area near a rigid polystyrene bead (large white circle, lower left). Phase contrast imaging; vertical white band is a camera artifact from the overexposed bead. White arrowhead: cell body initially had a thin, incomplete phase halo at t = 0 min; yellow arrowhead: at t = 3 min, the cell body was surrounded by a thick phase halo, suggesting increased height as compared to earlier. **(C)** Contours of the cell and bead shown in panel (B), registered to the stationary bead. As the cell moves closer to the bead, the cell adopts a classic unconfined cell shape. **(D)** Schematic of the PDMS confiner device. *(Right)* A suction cup lowers a piston in the presence of a vacuum. *(Left)* PDMS spacers hold the ceiling off the substrate (spacers are 3 µm tall and >1 mm apart). **(E)** Membrane-labeled keratocyte (CellMask Deep Red) migrating in the PDMS confiner. The ceiling was initially in the upright position, then was lowered gradually during the first 2 minutes of the time-lapse. (*Left*) Phase contrast images. (*Right*) Maximum-intensity projections into the indicated planes. XZ-plane projections are gamma-corrected to better visualize the entire cell. Cells in **(B,E)** were isolated from wild-type 2 dpf larvae.

To improve reproducibility, imaging, and temporal control of confinement, we implemented a vacuum-controlled confinement device that lowers a polydimethylsiloxane (PDMS) ceiling to a height of ∼3 µm (Le Berre et al., 2012). This flat, rigid, polyethylene glycol-coated ceiling also provides simpler geometry, mechanics, and surface chemistry compared to a viscoelastic agarose gel (**Figure 2D, S1**). When the PDMS ceiling is lowered onto isolated keratocytes, they display rapid shape changes. In XZ-plane (side view) projections, the cell body becomes flattened while the lamellipodium bulges upwards, often reaching the height of the ceiling. In the XY-plane (top-down) view, the lamellipodium becomes narrower, as the cell body expands in area, elongating the cell parallel to the direction of migration (**Figure 2E, Video S4**). Together, the agarose gel and PDMS ceiling experiments demonstrate that restricting the height of keratocytes induces elongated, narrower cells, irrespective of the confining material.

### Moderately confined keratocytes fall between unconfined and *in vivo* states on a height-length continuum

Combining reproducible confinement via the PDMS confiner and geometrically simple keratocytes enabled a quantitative study of the interplay between environment dimensionality and cell shape. Keratocytes were isolated from 2 dpf larvae stably expressing cytoplasmic mCherry and then labeled with the far-red membrane dye CellMask Deep Red. We made parallel measurements on keratocytes isolated from mCherry-negative sibling larvae to control for mCherry phototoxicity. Each cell was imaged in the PDMS confiner with the ceiling in the raised position to characterize both its three-dimensional shape and its motility behavior. Under our standard imaging conditions, collection of a set of high-resolution z-stacks (240 nm slices) for 10-15 fields of view in a single sample could be completed in about two minutes. After this initial shape determination, all cells in the sample were re-imaged using single slices at the level of the lamellipodium using two paired images at exactly 1 minute apart in order to accurately measure cell speed. Next, a second round of z-stack images were collected immediately before the ceiling was lowered. After lowering the ceiling and restoring proper focus, we continued imaging the same fields of view, alternating collection of z-stacks and one-minute single-slice speed measurements, for 20-30 minutes. Using this approach, we collected shape and speed data on large populations of the same individual cells before and after confinement (**Figure 3A, left**). Some cells did not have adherent lamellipodia and exhibited circular or irregular shapes, which we categorized as unpolarized (**Figure 3A, third column**). Other cells ruptured upon confinement, as evidenced by scattered puncta of previously cytoplasmic mCherry (**Figure 3A, fourth column**). All time frames were individually scored for these two behaviors (*Methods*). Unpolarized or ruptured states occurred spontaneously in 12% of unconfined cells, 41% of cells confined to a height greater than 2.5 µm, and 85% of cells confined to less than 2.5 µm (**Figure 3B**). Unpolarized cells were effectively non-motile compared to their polarized counterparts (**Figure S2A**). For further analyses, we retained only cells which were able to continue migrating with persistent polarity after confinement. Among this population, the cells had an average height of 8 µm for the unconfined condition and 4 µm for the confined condition (**Figure S2B**).

**Figure 3.**
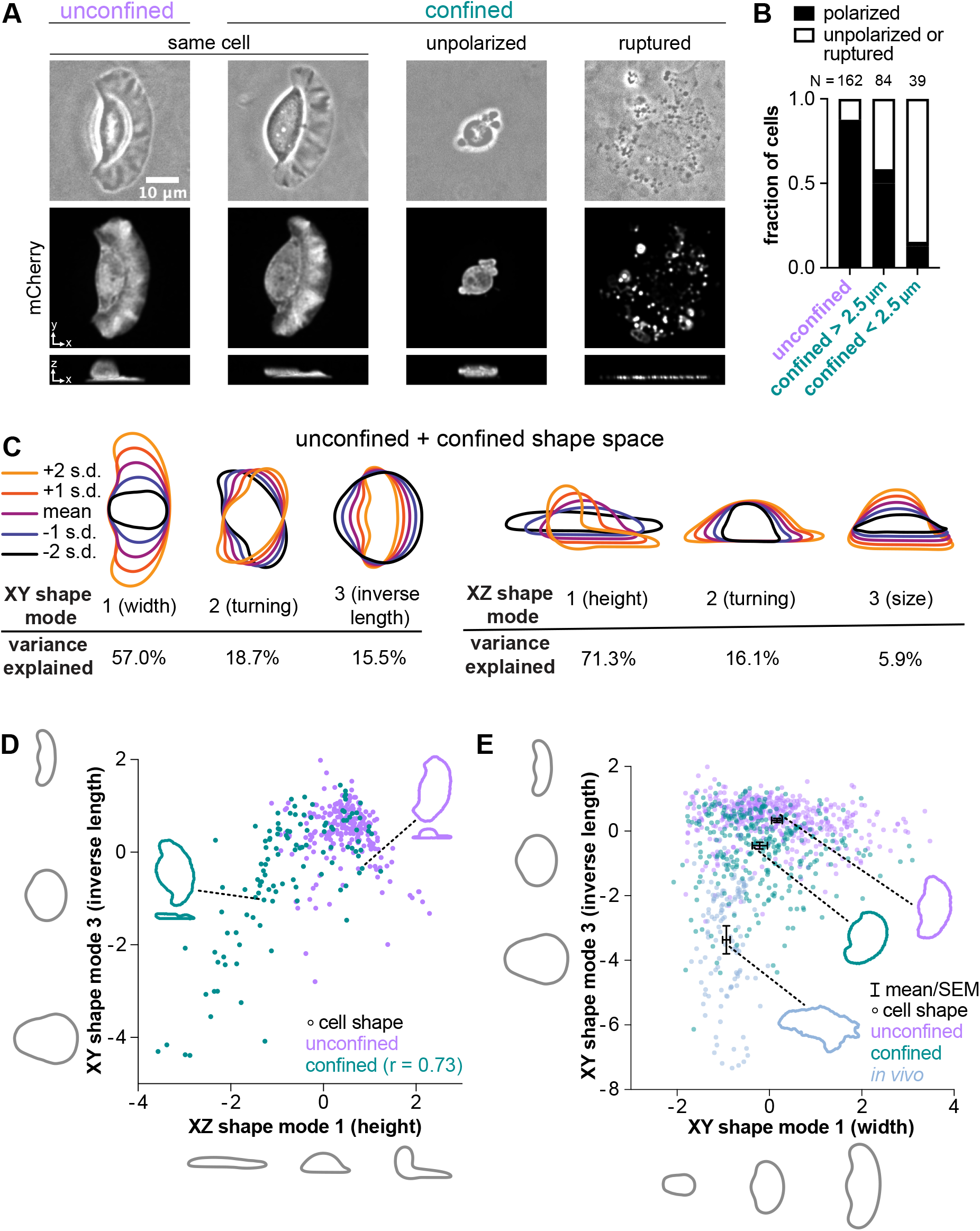
Moderately confined keratocytes fall between unconfined and *in vivo* states on a height-length continuum. **(A)** Representative images of keratocytes upon confinement. Keratocytes isolated from 2 dpf larvae expressing mCherry (*TgBAC(ΔNp63:Gal4)*; *Tg(UAS:mCherry))* were imaged in the PDMS confiner before the ceiling was lowered (left image) and during confinement (right three images). *(Top)* Phase contrast images. *(Bottom)* Maximum-intensity projections. **(B)** Portion of cells that displayed the behaviors illustrated in (A) before and during confinement (n ≥ 39 for all groups). Cells that displayed both polarized and non-polarized behavior over the course of the acquisition are categorized as polarized. **(C)** Modes of shape variation for the combined unconfined/confined keratocyte population, as determined by principal component analysis (PCA) of velocity-aligned outlines. *(Left)* The first three XY shape modes from PCA of 798 outlines of cells expressing mCherry or labeled with CellMask Deep Red. *(Right)* The first three XZ shape modes from PCA of 371 outlines of cells expressing mCherry. **(D)** Plot of all unconfined and confined coefficients along the first XZ and third XY shape modes. Idealized outlines along each axis were generated by varying that shape mode while holding the coefficients of the other modes at zero. The example outlines within the plot correspond to the cell from the left half of (A). Dots: observations of cell shape (before, n = 225; after, n = 119; with partial pairing and repeated cell measurements), from 8 samples. Effect sizes for all shape modes are shown in Table S1. **(E)** Plot that overlays 1) Unconfined and confined isolated cell shape coefficients, and 2) Shape coefficients of *in vivo* cells, measured by projecting their outlines into the shape space shown in panel (C). *In vivo* outlines were segmented from the first 10 minutes after wounding 3 dpf larvae mosaically expressing either a LifeAct marker or cytosolic and membrane markers (**Figure S3B**, *Methods*). Dots: observations of cell shape, including repeated cell measurements (n ≥ 135 observations for each condition). Example outlines within the plot: cells that have nearly average coefficients for their respective conditions along XY shape modes 1 and 3. Error bars: mean and standard error of the mean (SEM) for the condition indicated by the nearby example cell outline. Means and SEMs were calculated from 12 unconfined samples, 11 confined samples, and 4 larvae. Statistical comparisons are shown in Table S2.

This dataset allowed a quantitative examination of the effect of confinement on multiple shape variables. We used an unbiased, information-rich approach built on Principal Component Analysis, or PCA, for cell shape analysis (Pincus & Theriot, 2007). PCA requires that cell shapes be represented as vectors. To construct vectors that meaningfully represent cell shape, we segmented the CellMask or mCherry channel for each cell, aligned the mask to the direction of its instantaneous velocity, projected the mask into the XY or XZ plane, then extracted a set of point coordinates that outlined the cell boundary (**Figure S3A,C,** *Methods*). We conducted PCA separately for both the XY- and XZ-plane datasets, thereby determining the axes of greatest shape variation across the combined confined and unconfined populations. We refer to each axis as a *shape mode*, and the combined set of axes as a *shape space*. XZ-plane segmentations from mCherry-negative cells were not included in the PCA, due to heterogeneous CellMask signal near the apical surface (**Figure S3A**). For the XY-plane PCA, mCherry labeling did not have a significant effect on any of the top three shape mode scores (p > 0.05 for each mode, two-way ANOVA for label and confinement condition), so we pooled mCherry-positive and mCherry-negative samples.

In both the XY and XZ planes, just 3 shape modes describe over 90% of shape variation (**Figure 3C**). These shape modes can be assigned human-interpretable meaning by generating the outlines of idealized cells as they varied along each mode, while holding the coefficients of other modes constant at zero. For example, XZ shape mode 1 appears to be largely correlated with cell height (**Figure S2C**). Additionally, a cell with a positive XZ shape mode 1 coefficient has a round cell body that is much taller than the lamellipodium. A negative coefficient along XZ shape mode 1 indicates a thin cell with uniform height (**Figure S2D**). This shape mode describes 71% of the XZ shape variation. The average value of the coefficient for XZ shape mode 1 was lower for confined cells as compared to unconfined ones (**Table S1**), confirming our initial observation that confinement induced unusually flat cells, as expected given the mode of confinement.

The axes that best distinguish confined and unconfined cell shapes are XZ shape mode 1 and XY shape mode 3, which appears to primarily report on the cell’s length parallel to the direction of migration (**Figure S2E**). For both modes, the confined cells have exaggerated negative scores, indicating flat, highly elongated cells (**Figure 3D**). In contrast, other shape modes showed much smaller effect sizes between confined and unconfined cells (**Table S1, Figure S2F**). There was also a strong correlation between height and inverse length for confined cells (Pearson r = 0.73, p < 0.0001). Therefore, cell elongation in the XY plane occurs concomitantly with Z-plane confinement, even though the confinement method does not impose any boundaries in the XY plane. Elongation may occur simply because the cell body and the dense lamellipodium cannot be stacked vertically, and instead the cell body is carried behind the lamellipodium for confined cells, while it is carried above the rear axle of the lamellipodium in unconfined cells. Alternatively, the confined cell could be longer due to differential friction on the cell body versus the lamellipodium. Regardless of the mechanism of shape change, elongation in the direction of migration is a stereotypic response to confinement.

Next, we used this PCA-generated shape space, developed on measurements of isolated cells only, to map the differences between *in vivo* and isolated keratocyte shapes. We measured *in vivo* shapes using 3 dpf larvae mosaically expressing either LifeAct-mNeonGreen or cytoplasmic mNeonGreen with a membrane-tagged mRuby. Cells were imaged during the first 10 minutes after laceration and manually segmented in the XY plane using both channels (**Figure S3B**). We extracted only the XY shape because the XZ shape was obscured by the tailfin’s slight curvature and tilt relative to the imaging plane. These cells were projected into the isolated cell shape space, with the *in vivo* wound direction aligned to the isolated cell velocity direction. Despite large cell-to-cell variability, *in vivo* and unconfined cells were well separated in the shape space, with *in vivo* cells scoring as narrower and longer than the isolated cells (**Figure 3E, Table S2**). In addition, *in vivo* cells are more variable in length than width, whereas unconfined cells are more variable in width than length. Confined cells are variable along both modes, with individual scores overlapping both unconfined and *in vivo* scores. The average shape coefficients for all three populations are surprisingly collinear, with confined cells falling between unconfined and *in vivo*. Therefore, 2D confinement phenocopies *in vivo* shape by elongating cells in the direction of migration, although confinement does not fully mimic the narrowness or lamellipodial ruffling of *in vivo* cells. These shape features may be a product of cell-cell interactions or mechanical properties not recapitulated by the rigid PDMS ceiling. *In vivo* shape similarity is best achieved by moderate confinement that substantially deforms the otherwise spherical cell body, without rupturing the cell. These quantitative comparisons demonstrate that 2D confinement makes a significant contribution to cell shape in the larval zebrafish epidermis.

### Confining basal cells reduces speed and volume

Our paired image data additionally enabled an investigation into the effect of confinement on keratocyte speed, to understand how the skin environment may or may not promote cell migration during wound healing. In the pairwise comparison before and during confinement, isolated keratocytes showed a speed decrease of 22% ± 8% (s.d., n=10 samples, **Figure 4A**). As a control, keratocytes were measured using the same imaging protocol without confinement. Under these conditions, speed showed no overall change over a similar duration (**Figure S4A, left**). To explore possible influences on speed, we reconstructed the cell boundary in three dimensions (**Figure 4B**). Surprisingly, this analysis revealed confinement decreased cell volume by 21% ± 1% (s.d., n=5 samples, **Figure 4C**). In contrast, unconfined control samples maintained volume within 2% ± 3% (s.d., n=4 samples, **Figure S4A, middle**). Surface area measurements were not significantly different between unconfined and confined conditions (**Figure 4D**). This observation is consistent with previous keratocyte surface electron micrographs and tether pulling measurements, which indicated that keratocytes maintain a fully extended plasma membrane under high tension (Lieber et al., 2013). These results suggest that in response to extreme cell body deformation, keratocytes lose volume overall because they can neither expand their membrane area nor redistribute sufficient volume from the cell body to the lamellipodium.

**Figure 4.**
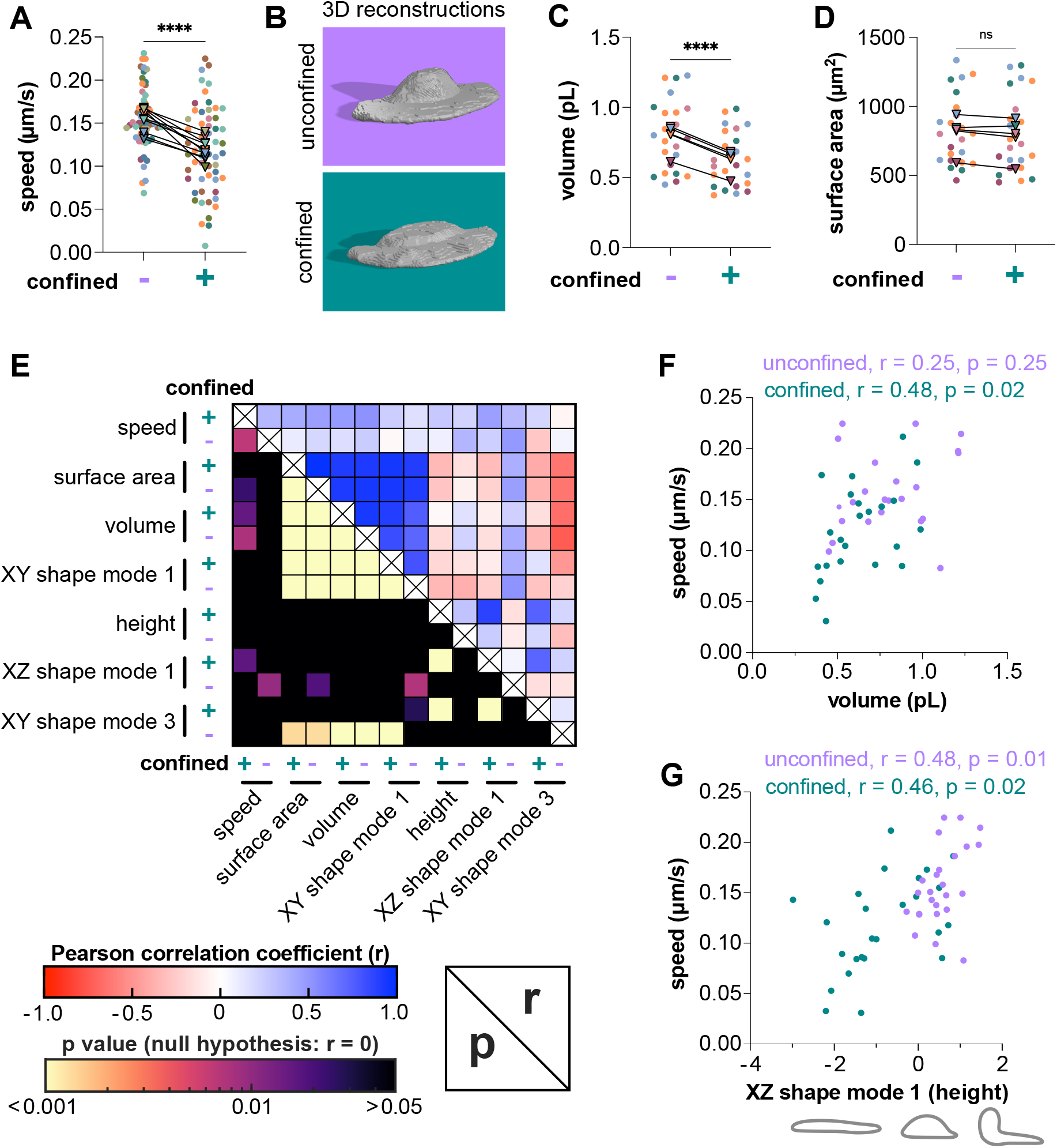
Confining basal cells reduces speed and volume. **(A)** Pairwise speed comparison of cells before and during confinement in the PDMS confiner, from the same dataset shown in Figure 3 (pooling mCherry-positive and negative samples). Centroid speed was measured over a 60-second interval, 5-10 minutes before confinement (“unconfined”) and again 6-10 minutes after the ceiling was lowered (“confined”). Paired t-test conducted on 10 sample averages (n ≥ 3 cells from each sample), with p < 0.0001. For **(A, C, D)**, dots indicate individual cell measurements and triangles indicate sample averages. The lines connect the same sample across conditions. Cell and sample values are color-coordinated, with each color representing a different sample. **(B)** Rendered 3D meshes of the example cell from Figure 3A in its unconfined and confined states. Meshes were reconstructed from deconvolved confocal images of cytoplasmic mCherry. **(C)** Pairwise volume comparison before and during confinement, measured using 3D reconstructions as shown in (B). Unconfined measurements were taken twice 3-15 minutes before confinement and averaged. Confined measurements were taken 4-8 minutes after the ceiling was lowered. Paired t-test conducted on 5 sample averages (n ≥ 3 cells from each sample), with p < 0.0001. **(D)** Pairwise surface area comparison before and during confinement, measured and sampled as in (C). Paired t-test, with p > 0.05 (ns: not significant). **(E)** Correlation matrix of all possible pairs of speed, size, and shape measurements before and during confinement. Pairwise Pearson correlation coefficients (r) were calculated on individual cells, which were each measured both with and without confinement (n ≥ 23). Correlation coefficients are shown on the top-right of the diagonal, and p-values from two-sided t-tests are shown to the bottom-left of the diagonal. **(F)** Plot of volume versus speed for confined and unconfined measurements. Each dot represents an individual cell (n = 23, with each cell measured in both conditions). For **(F,G)**, Pearson correlation coefficients (r) and p-values are calculated as in panel E. **(G)** Plot of XZ shape mode 1 versus speed for confined and unconfined measurements. Each dot represents an individual cell (n = 25, with each cell measured in both conditions).

Cell size, shape, and speed changes were measured within the same paired dataset, which allowed us to explore the relationships among variables. Pearson correlation coefficients were calculated between all possible pairs of measurements for both unconfined and confined cells on a per-cell basis and the results summarized as a heat map (**Figure 4E**). Surface area, volume, and XY shape mode 1 (width) formed a cluster of cell size variables that were all highly correlated with one another across confinement conditions. The variables that define the flattening/elongation response, including height, XZ shape mode 1 (height), and XY shape mode 3 (inverse length), showed high correlations between confined shape values and low correlations between unconfined values, indicating that confined shape is unrelated to prior cell shape properties. Overall, the heat map emphasized strong correlations between many size and shape variables.

To generate hypotheses about causes of cell speed loss under confinement, we investigated correlations involving both confined and unconfined speed measurements (**Figure 4E**). Notably, confined volume and confined speed exhibited a correlation coefficient of 0.48 (**Figure 4F**). Confined speed correlated similarly with unconfined volume, unconfined surface area, and confined surface area, which was expected given the strong correlations within the cell size cluster of the heat map. In contrast, size variables were not correlated with unconfined speed, suggesting that the size/speed relationship is specific to the application of confinement. XZ shape mode 1 (height), although uncorrelated with the cell size cluster, also showed significant correlations with speed (**Figure 4G**). The correlation coefficient was 0.48 before confinement and 0.46 during confinement, indicating that height is associated with speed independently of applied environmental changes. Finally, cell-to-cell speed variability persisted across conditions, indicated by a good correlation between unconfined and confined speed (**Figure S4B**). Therefore, cell size, height, and cell-intrinsic properties are all indicative of cell speed immediately after confinement.

### Hypertonic shock causes volume and speed loss on par with confinement

Cell volume changes in response to confinement are likely to affect cytoplasmic properties, and by extension, cytoskeletal dynamics. In order to directly test a potential volume-speed relationship, we altered keratocyte volume in a manner unrelated to physical confinement. Specifically, we used hypertonic shock to rapidly decrease keratocyte volume. We made alternating 3D shape and speed measurements for up to 35 minutes using our standard imaging protocol. During the acquisition, the medium was swapped for media supplemented with 20-50 mg/ml sorbitol, which does not enter animal cells (Watari et al., 2004; Wood et al., 1968). The hypertonically-shocked cells maintained lamellipodial migration, displaying the standard unconfined keratocyte shape with a large fan-shaped lamellipodium (**Figure S4C**). Pooling across sorbitol concentrations, the hypertonic treatments achieved volume changes comparable to those observed in confinement (**Figure 5A,B**). Surface area remained constant during the volume decrease, again suggesting that keratocytes do not remodel their membrane on the timescale of minutes (**Figure 5C**). Interestingly, hypertonic shock slowed cells down by 20% ± 10% (s.d., n=11 samples, **Figure 5D**), similar to the speed decrease we had observed for confined cells.

**Figure 5.**
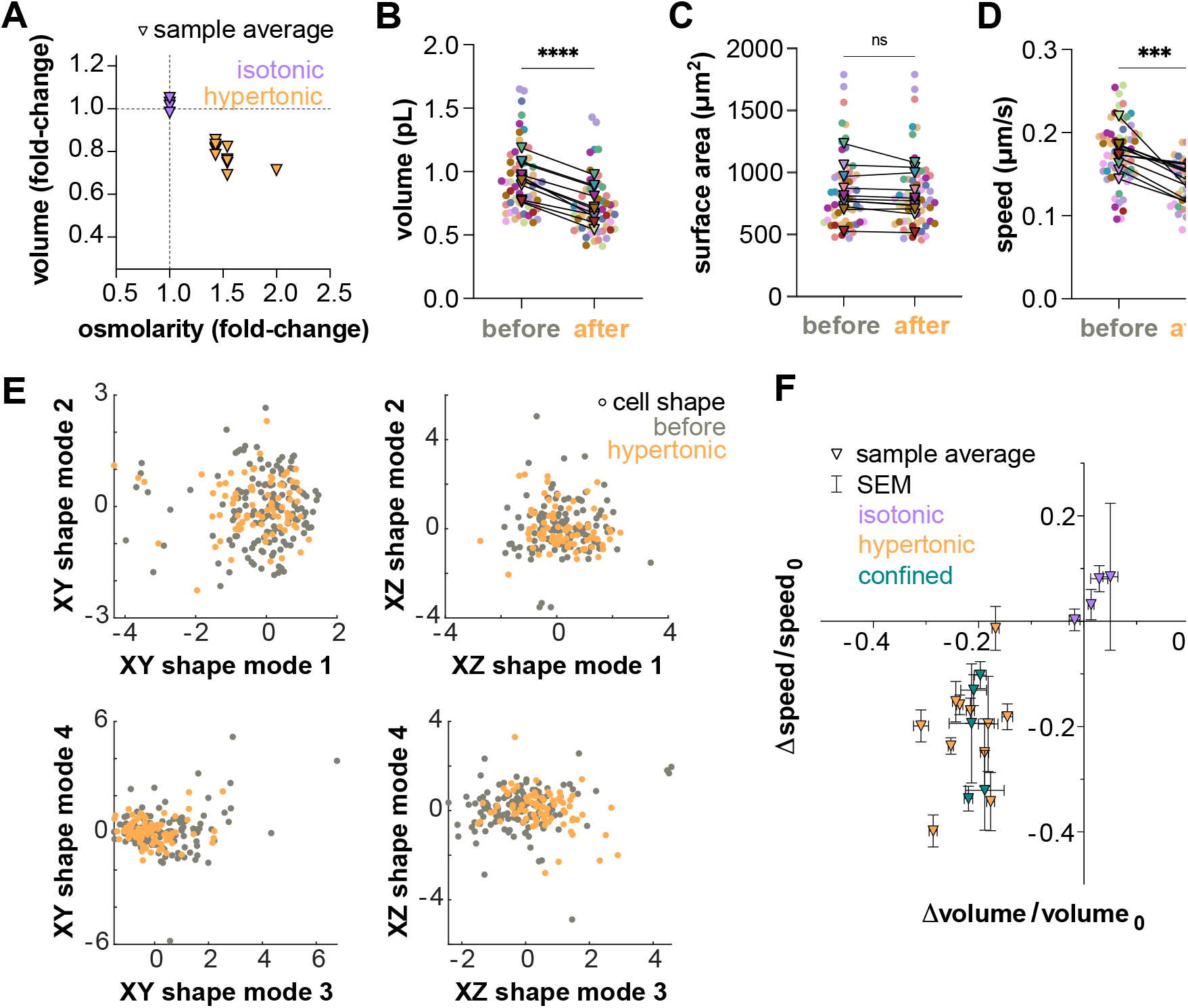
Hypertonic shock causes volume and speed loss on par with confinement. **(A)** Keratocytes isolated from 2 dpf larvae expressing mCherry were imaged during a media shift to either the original isotonic medium or media supplemented with 20-50 mg/ml sorbitol. Plot indicates sample-averaged fold-changes, with osmolarity estimated from media and sorbitol concentrations. **(B)** Detailed volume comparison of cells before and after hypertonic shock. Volume was measured using 3D reconstructions of cytoplasmic mCherry. Volumes were averaged between 2-8 minutes preceding the media swap (“before”), and from immediately after the media swap until cells left the field of view or 30 minutes had elapsed (“after”). Paired t-test conducted on 11 sample averages (n ≥ 3 cells from each sample), with p < 0.0001. For **(B-D)**, dots indicate individual cell measurements and triangles indicate sample averages. The lines connect the same sample across conditions. Cell and sample values are color-coordinated, with each color representing a different sample. **(C)** Pairwise surface area comparison of cells before and after hypertonic shock, measured and sampled as in (B). Paired t-test, with p > 0.05 (ns: not significant). **(D)** Pairwise speed comparison of cells before and after hypertonic shock. Centroid speed was measured over 60-second intervals, spaced out by longer z-stack acquisitions. Before speeds were averaged within 6 minutes preceding the media swap. After speeds were averaged from 3 minutes after the media swap until cells left the field of view or 30 minutes had elapsed. Paired t-test conducted on 11 sample averages (n ≥ 3 cells from each sample), with p = 0.0002. **(E)** Plot of shape coefficients before and after hypertonic shock, from the top four shape modes in the XY plane (left half) or XZ plane (right half), showing little change between before and after. Effect sizes for top six shape modes are shown in Table S3. Idealized contours for each shape mode are shown in Figure S4D,E. Dots: average shape for each cell during observation before and after hypertonic shock (before, n = 145; after, n = 83; with partial pairing), from 14 samples. **(F)** Plot that summarizes the volume and speed changes observed for samples placed under confinement, switched to hypertonic media (supplemented with 20-50 mg/ml sorbitol), or switched to isotonic media (same data shown in Figure 4A,C, Figure 5B,D, and Figure S4A). Differences were calculated for individual cells, normalized to the initial value, then averaged (N ≥ 4 samples per condition). Error bars indicate the standard error of the mean

To extend our observations on the effect of hypertonic shock to other aspects of keratocyte shape in addition to total cell volume, we generated a second shape space (using the methodology described above) combining cells after treatment with hypertonic medium and control cells in isotonic medium (**Figure S4D,E**). The principal shape modes in the XY plane were strikingly similar to the modes established for the combined unconfined and confined cell populations (compare **Figure S4D** and **Figure 3C, left half**). However, the modes in the XZ plane were qualitatively different, reflecting the observation that cells treated with hypertonic shock never achieved the completely flat profiles of confined cells (compare **Figure S4E** and **Figure 3C, right half**). Within this new shape space, we compared individual cell shapes before and after hypertonic shock. Overall, there was very little change in shape mode coefficients for hypertonically-shocked cell shapes versus their pre-shock values (**Figure 5E**). XZ shape mode 3 had the biggest difference between conditions, with an effect size of 1.5. However, this mode was highly correlated with cell volume and therefore does not describe a distinct shape change separate from the volume-dependent effect (**Figure S4F**). If XZ shape mode 3 is not considered, the shape mode effect sizes for hypertonic treatment are all within ±0.65 standard deviations, which is much smaller than the effect sizes for confinement of >2 standard deviations (**Table S1,S3**). Therefore, we conclude that only a change in volume, and not a volume-independent change in shape, is associated with speed loss following hypertonic shock.

Keratocytes responded to confinement and hypertonic shock with comparable losses in volume and speed, despite dramatic differences in type of perturbation and resulting cell shape (**Figure 5F**). In both conditions, cells lost around 20% of both their speed and volume, with a similar level of sample-to-sample variability in either experiment. This suggests that speed reduction from both confinement and hypertonic shock may involve a shared mechanism, mediated through rapid volume loss.

## Discussion

Here, we examined keratocytes *in vivo*, in culture, and under artificial 2D confinement to evaluate the effect of skin-like geometric constraints on lamellipodial migration. We demonstrated that the larval skin environment deforms *in vivo* keratocytes relative to their isolated shape, so that they are flattened in height and elongated parallel to the direction of migration. A similar effect may occur in the multi-layered adult zebrafish epidermis, which also displays lamellipodial migration with cells elongated substantially in the direction of migration (Richardson et al., 2016). 2D confinement in culture is sufficient to recapitulate this physical constraint and its effect on cell shape, without need of neighboring cells or specific adhesions. However, despite having a more physiological shape, confined keratocytes are slower than their unconfined counterparts.

It is possible that shape changes partially contribute to confined speed loss. Our cross-sectional shape analysis revealed that cell roundness is connected to cell speed. As measured with XZ shape mode 1, we observed that flat, thin cells tended to be slower than those with tall, spherical cell bodies both before and during confinement (**Figure 4G**). A possible explanation for this correlation is that a round cell body may promote faster cell speed, consistent with a previously proposed “cell body rolling” mechanism of translocation of keratocytes (Anderson et al., 1996; Okimura et al., 2018). The rolling model proposes that the rotation of cell body material acts like a wheel rolling in the direction of migration, partially contributing to cell speed. It seems likely that a confined cell body, which has a much less circular XZ cross-section, would roll less efficiently. Cell body rotational tracking, in parallel with shape analysis, could clarify whether altered rolling rates explain the relationship between roundness and speed.

The hypertonic shock experiments suggest that keratocyte speed is partially determined by cell volume or a physical property correlated with volume, such as cytoplasmic density. A 20% reduction in volume from either hypertonic shock or confinement was associated with an equivalent percent reduction in speed (**Figure 5F**), even though cells were more deformed under confinement than hypertonic shock. This substantial volume change during active migration is surprising, given that the cell is only 60-80% water and that many biochemical and diffusive processes are sensitive to protein concentration and cytoplasmic macromolecular crowding (Milo & Phillips, 2016; Molines et al., 2022; Neurohr & Amon, 2020). Nonetheless, similar volume-speed trends have been reported for 1) primary human neutrophils in various media tonicities (Rosengren et al., 1994), 2) cancer cells in 3D hydrogels of various stiffnesses (Wang et al., 2020) and 3) *Dictyostelium* amoebae under 2D confinement with various applied loads (Srivastava et al., 2020). All three studies reported that conditions yielding smaller cell volume also caused slower migration speeds, in agreement with our findings. Using paired single-cell measurements and quantitative shape analysis, this work more directly connects speed and volume, by decoupling volume changes from geometric perturbations and other alterations in cell shape.

The underlying connection between speed and volume remains unidentified. Based on a lack of cell rupture or vesiculation, it seems likely that cells lose water volume and therefore experience a change of cytoplasmic physical properties. Rapid water flux through aquaporins has been demonstrated many times in response to osmotic shock (Preston et al., 1992). Recently, volume loss due to 2D confinement was measured to also occur rapidly, with HeLa cells losing 30% of cell volume within 30 milliseconds (Venkova et al., 2022). This volume flux is a reasonable rate for ∼100,000 aquaporin channels (Gade & Robinson, 2006). Thus, water efflux is a likely cause of confinement-induced volume loss.

Water loss leads to substantial cytoplasmic changes, as measured by increases in cell stiffness and decreases in diffusion rates (Guo et al., 2017; Joyner et al., 2016; Miermont et al., 2013; Zhou et al., 2009). These changes occur independently of an intact actin cytoskeleton and therefore are thought to operate via general cytoplasmic molecular interactions. Interestingly, osmotic-induced water flux has been recently shown to have a dramatic effect on microtubule polymerization and depolymerization in fission yeast (Molines et al., 2022). Here, volume loss, rather than accelerating polymerization via an increase in tubulin concentration, instead slowed tubulin diffusion and ultimately polymerization and depolymerization. Actin polymerization is a diffusion-limited reaction and therefore likely to be sensitive to cellular properties that affect cytoplasmic diffusion rate, like volume (Drenckhahn & Pollard, 1986).

Additionally, keratocytes require global actin transport because network assembly is high at the front of the cell, while disassembly dominates at the rear (Wilson et al., 2010). Normally, actin monomers can diffuse across the cell in 3 seconds and are further aided by bulk fluid flow from back to front (Keren et al., 2009; Raz-Ben Aroush et al., 2017). Water efflux could perturb these transport processes and thereby additionally reduce actin availability for polymerization at the cell front. Therefore, loss of cytoplasmic water may impede keratocyte migration via altered actin diffusion and transport, while leaving cytoskeletal shape and organization relatively intact. In tissues, osmotic shock is reported to affect wound closure rates across a number of species. Axolotl skin explants heal slower in the presence of hypertonic media (Tanner et al., 2009). With the opposite perturbation, hypotonic shock, zebrafish larvae and *Drosophila* embryos heal faster (Gault et al., 2014; Kennard & Theriot, 2020; Scepanovic et al., 2021). Zebrafish, as a freshwater species, are thought to use such hypotonic shock as a normal signal for migration after wounding. It is tempting to speculate that hypotonic shock promotes wound healing not just through signaling pathways, but also through physical effects on cells. Indeed, individual keratocytes swell to 50% of their pre-wounding volume following normal wounding in freshwater (Kennard & Theriot, 2020), which could significantly boost speed if the volume-speed relationship holds in the swelling regime. Hypotonic shock additionally leads to a massive fluid influx that increases the space between basal cells and the overlying periderm (Kennard et al., 2022). This expansion of the epidermis may release some of the 2D constraints on keratocytes and provide additional speed benefits. The evidence presented here suggests that hypotonic shock may promote wound closure by physically increasing cell migration speed, potentially by both diluting the crowded cytoplasm and reducing 2D confinement.

## Acknowledgments

We are grateful to Christopher Prinz for drawing the schematic in Figure 1C and Andrew Kennard for sharing fish lines and plasmids, as well as advice concerning many aspects of this project. We are also grateful to Emily Hatch for sharing her expertise in confinement methods. We thank Jeff Rasmussen and Alvaro Sagasti for sharing fish lines, Philippe Mourrain and David Raible for hosting fish in their facility, as well as David White and George Sanders for their fish husbandry expertise. We are grateful to Ramon Lorenzo D. Labitigan for sharing code and expertise on shape analysis. Finally, we thank Kinneret Keren and Aaron van Loon for careful critical reading of the manuscript and the members of the Theriot laboratory for their insightful discussions and feedback.

Funding: Support for JAT comes from Howard Hughes Medical Institute and the Washington Research Foundation. Support for MJF comes from Howard Hughes Medical Institute. Support for ECL comes from Howard Hughes Medical Institute and the National Science Foundation Graduate Research Fellowship Program under Grant No. DGE-1656518. Any opinions, findings, and conclusions or recommendations expressed in this material are those of the authors and do not necessarily reflect the views of the National Science Foundation. Part of this work was conducted at the Washington Nanofabrication Facility, a National Nanotechnology Coordinated Infrastructure (NNCI) site at the University of Washington with partial support from the National Science Foundation via awards NNCI-1542101 and NNCI-2025489.

This article is subject to HHMI’s Open Access to Publications policy. HHMI lab heads have previously granted a nonexclusive CC BY 4.0 license to the public and a sublicensable license to HHMI in their research articles. Pursuant to those licenses, the author-accepted manuscript of this article can be made freely available under a CC BY 4.0 license immediately upon publication.

## Author contributions

Ellen C. Labuz: conceptualization, investigation, methodology, software, formal analysis, visualization, writing – original draft. Matthew J. Footer: methodology, resources, visualization, writing – review & editing. Julie A. Theriot: conceptualization, supervision, funding acquisition, writing – review & editing.

## Materials and Methods

### Zebrafish husbandry

Zebrafish (TAB5 background wildtype strain) were maintained according to standard procedures (Westerfield, 2007). Experiments were approved by University of Washington Institutional Animal Care and Use Committee (protocol 4427-01). Animals were raised on a 12 hr light, 10 hr dark cycle at 28.5 °C. Adults were crossed through natural spawning. Embryos were collected and raised in 100 mm petri dishes at 28.5 °C. Each dish contained 30-50 embryos in system water.

### Transgenic zebrafish lines

The following previously generated transgenic zebrafish lines were used: *TgBAC(ΔNp63:Gal4)^la213^* (Rasmussen et al., 2015), *Tg(UAS:LifeAct-EGFP)^mu271^* (Helker et al., 2013), *Tg(UAS:mCherry)* (Kennard et al., 2022).

### Plasmid constructs and mRNA synthesis

The UAS:mNeonGreen-P2A-mRuby3-CAAX as previously published (Kennard & Theriot, 2020). The UAS:LifeAct-mNeonGreen-P2A-mRuby3-CAAX plasmid was generated using the Tol2kit (Kwan et al., 2007) using standard Gateway cloning methods (Invitrogen). Briefly, the final plasmid was formed by LR reaction (LR Clonase II, Invitrogen) to recombine plasmids p5E-UAS (Tol2kit), pME-LifeAct-mNeonGreen-P2A (this work), and p3E-mRuby3-CAAX (Kennard & Theriot, 2020) into destination vector pDestTol2CG2 (Tol2kit), containing the cmlc2:GFP transgenic marker. The construction of the p3E-mRuby-CAAX plasmid has been previously published (Kennard & Theriot, 2020). The pME-LifeAct-mNeonGreen-P2A plasmid was generated from the previously published pME-mNeonGreen-P2A (Kennard & Theriot, 2020) using Q5 mutagenesis with primers #1 and #2 (**Table S4**) to insert the LifeAct peptide (Riedl et al., 2008) in-frame at the N terminus. The pME and p3E insert sequences were confirmed by Sanger sequencing and the final destination vector was confirmed by restriction digest.

Tol2 transposase mRNA was synthesized using the SP6 mMESSAGE mMACHINE reverse transcription kit (Invitrogen), with the Tol2kit plasmid pCS2A-transposase as a template. All Tol2kit plasmids were a gift from C.-B. Chien.

### Microinjection for mosaic expression

Embryos from the *TgBAC(ΔNp63:Gal4)^la213^* line (Rasmussen et al., 2015) were injected at the 1- to 2-cell stage. The injection mix contained 20 ng/µl of the UAS-driven plasmid and 40 ng/µl of Tol2 mRNA. The volume of the injected drop was not calibrated but was approximately 2 nl to achieve mosaicism. At 3 days post fertilization (dpf), larvae were screened for mosaic expression, with approximately ∼5% of tailfin basal cells expressing the UAS-driven fluorescent protein.

### Tissue laceration

Experiments were performed at 3 dpf. Mosaically-expressing larvae were prepared for imaging and lacerated as previously described (Kennard et al., 2021). Briefly, larvae were anesthetized in larval imaging media, a mixture of E3 (5 mM NaCl, 0.17 mM KCl, 0.33 mM CaCl2, 0.33 mM MgSO4), 160 mg/ml Tricaine (Sigma), and 0.8 mM Tris pH 7. Larvae were mounted in 2% agarose (Invitrogen) onto a 25 mm coverslip. Larvae were mounted on their right side, with the dorsal-ventral axis aligned parallel to the coverslip. The cooled agarose was covered with excess media and a portion of the agarose around the tail fin was removed. Before wounding, a 60X magnification field of view, which included multiple construct-expressing cells, was selected from the tail fin either dorsal or ventral to the notochord. Laceration was performed with needles pulled from solid borosilicate glass rods (Sutter BR-100-10) using a Brown-Flaming type micropipette puller (Sutter P-87). The needle was dragged through the tailfin twice, from near the end of the notochord towards the posterior edge of the tailfin at a 45° angle.

### Keratocyte isolation and cell culture reagents

Keratocytes were isolated from 2 dpf larvae, which were either wild-type or screened at 1-2 dpf for transgenes of interest. Isolation was performed as previously described (Lou et al., 2015). Briefly, larvae were dechorionated and anesthetized with 160 mg/ml Tricaine. Larvae were washed twice in PBS, incubated in cell dissociation buffer (Fisher) at 4°C for 30 minutes, mechanically mashed by a 200 µL pipette tip, and incubated in 25% trypsin and 1 mM EDTA for ∼15 minutes at 28°C. The trypsin was quenched with fetal bovine serum supplemented with antibiotic/antimycotic and the supernatant further concentrated by centrifugation for 3 minutes at 500 g. This cell-rich solution was plated on coverslips coated with rat tail collagen I (Gibco) and incubated for 1 hr at room temperature to allow keratocytes to adhere. The samples were washed three times with Leibovitz’s Media (L-15), supplemented with 10% fetal bovine serum and antibiotic/antimycotic, then imaged in the same media. Some samples were incubated at 4°C for 1-2 hours to reduce microbial growth, followed by 1 hour at room temperature before imaging. Membrane staining was performed immediately before imaging each coverslip, using CellMask Deep Red (Fisher) for 10 minutes. For hypertonic experiments, the imaging media was supplemented with 20-50 mg/ml D-sorbitol (Sigma) and filter-sterilized.

### 2D confinement by agarose gel overlay

On the day of each experiment, a 2X solution of 2% agarose (w/w, UltraPure Low Melting Point Agarose Powder, Invitrogen) was dissolved by heating and then kept fluid at 42°C. A 2X imaging solution was prepared by mixing 20% fetal bovine serum into L-15. The final solution of 1% agarose and 10% fetal bovine serum was mixed from the 2X agarose solution and 2X imaging media in a 1:1 (v/v) ratio. An agarose slab was made by pipetting the final agarose mixture between two microscope slides with four #1.5 coverslip spacers and allowed to gel at room temperature. Polystyrene beads (25 µm diameter, Polysciences 07313) were diluted 1:100 in phosphate saline buffer and 10 µl of the dilution was distributed on a sample of keratocytes isolated from larvae 2 dpf (see *Keratocyte isolation and cell culture reagents*). The agarose slab was carefully draped over the keratocyte and bead sample and allowed to settle for 10 minutes before imaging.

### 2D confinement by vacuum-controlled device

The vacuum-controlled confiner device was manufactured similarly to previous descriptions and comprised a polydimethylsiloxane (PDMS) suction cup with a piston, which was capped by a coverslip with a micropatterned PDMS surface (Le Berre et al., 2012, 2014). Device production required two molds, one for the suction cup and one for the micropatterned ceiling (**Figure 2D, S1**). The suction cup mold was designed for use on top of a 25 mm coverslip and consists of 6 separate parts. The stainless steel and aluminum parts were machined by the UW Physics Department Machine Shop (for exact drawings, see **Figure S5**). We used Autodesk Fusion 360 for design and rendering. The O-ring (#117) and glass disk (1” x 0.25”) are off-the-shelf parts (McMaster-Carr). The ceiling mold was manufactured from SU-8 photoresist by the Washington Nanofabrication Facility. A photomask protected pillars of 440 µm diameter, spaced in an array with 2.5 mm by 0.7 mm center-to-center spacing (**Figure S1C**). The final mold height for the pillars was 3.2 µm.

To manufacture each suction cup, Sylgard 184 (Thermo Fisher) was thoroughly mixed in a PDMS/cross-linker ratio of 10:1 (w/w) and degassed in a vacuum chamber to remove bubbles. All parts of the mold were rinsed with 5% Pluronic F127 before assembly. All parts of the mold, minus the glass disk, are pre-assembled with the inner ring merely sitting on the platform (see SI). The PDMS was poured into the mold to just over the height of the ledge that the glass disk rests on and degassed again. The glass disk was lowered into place and the assembly was baked at 80°C for 1 hr and allowed to cool. The suction cup was de-molded with the aid of 100% isopropanol and pierced with a 0.5-mm biopsy punch to create an inlet for a vacuum line. Alternative mold designs have been described previously, including an assembly of aluminum rings, coverslips, and glass slides (Le Berre et al., 2014).

The confinement chips were manufactured as previously described (Le Berre et al., 2014). Briefly, Sylgard 184 was thoroughly mixed in a PDMS/cross-linker ratio of 8:1 (w/w), degassed, and poured onto the micropatterned SU-8 mold (which was encased by aluminum foil to prevent PDMS from flowing off the flat mold). After further degassing, plasma-cleaned 8-mm coverslips were placed in the PDMS and pressed onto the patterned surface. The mold was baked on a preheated 95°C hot plate for 15 minutes, and coverslips were de-molded using 100% isopropanol and a razor blade at a flat angle, nearly parallel to the coverslip to avoid damaging the mold.

On the day of each experiment, each confinement chip was plasma-cleaned and incubated in 0.5 mg/ml pLL-PEG (Susos) for 1 hr, then equilibrated with cell culture media for 1 hr. The suction cup was sanitized with 70% ethanol and allowed to dry. The glass side of the confinement chip was placed onto the piston, where it self-adhered. Then, the suction cup was sealed onto the sample on the microscope with −8.5 kPa gauge pressure. This vacuum level held the ceiling >50 µm above the cells. To apply confinement, the vacuum was slowly increased, with periodic measurements of cell height using the membrane channel. The cells were considered confined at <4 µm, which required between −17 to −30 kPa, depending on the sample. Afterwards, suction cups were cleaned with dish soap and stored for future use; confinement chips were discarded.

### Microscopy and image acquisition

Imaging for the agarose confinement experiment was performed on a Nikon Ti-E inverted microscope with transmitted light using phase contrast, a 100x objective (NA 1.45, Nikon) with 1.5x intermediate magnifier. Images were acquired onto a Andor iXon EMCCD camera using µ-Manager (Edelstein et al., 2014). All other experiments were imaged using a spinning disk confocal setup, consisting of a Nikon Ti2 inverted microscope, a piezo-z stage (Applied Scientific Instruments PZ-2300-XY-FT), a Yokogawa CSU-W1 spinning-disk confocal with Borealis attachment (Andor), and a back-thinned EMCCD camera in 16-bit imaging mode (Andor DU888 iXon Ultra). Confocal illumination was supplied by a laser launch (Vortran VersaLase) with 50 mW 488 nm, 50 mW 561 nm, and 110 mW 642 nm diode lasers (Vortran Stradus). Either a 100X oil objective (NA 1.45, Nikon) or a 60X oil objective (NA 1.40, Nikon) were used, with a 405/488/561/640/755 penta-band dichroic (Andor) and single-band GFP, RFP, and far-red filters (Chroma 535/50m, 595/50m, and 700/75m respectively). Images were acquired at room temperature (20-23°C) using µ-Manager. For the experiments that tracked shape, size, and speed, 10-15 positions were imaged in one acquisition from each sample. The frame interval was set to 60 seconds, with z-stacks only imaged every 3 frames with a z-slice interval of 240 nm. Acquiring the full z-stack at each position actually took 1-3 minutes, but the frame interval between two subsequent non-stack frames was always 60 seconds (useful for accurate speed quantification, see *Cell trajectory analysis*). During vacuum-controlled confinement experiments, the pressure decrease caused a significant change in the z-position of the coverslip, so the focal plane of each position was reset between confinement application and further imaging.

### Cell trajectory analysis

Cells were tracked based on single-slice images of the membrane channel, taken near the coverslip. Tracks were calculated using custom code written in Matlab R2018b (available upon request). Briefly, cell trajectories were determined using the centroid of the segmented membrane signal. Segmentation was performed on the entire image for the first frame, which required manual gamma adjustment and thresholding. For subsequent frames, segmentation was automatically performed using Otsu thresholding within a 150 x 150 or 200 x 200-pixel region around the previous centroid. Subsequent positions for the same cell were assigned by calculating all possible cell-to-cell displacements between consecutive time points and matching cells through the minimization of total displacement across cells. The cell density from the isolation procedure was low enough that individual tracks could be identified with this approach, followed by manual curation to remove misidentified tracks. For the confinement experiments, a significant time gap occurred between the before-confinement acquisition and the during-confinement acquisition. Therefore, all matching tracks were manually assigned, based on the expected direction of cell movement, approximate cell size, and intracellular components visible in the phase contrast channel. Cell speeds were only calculated between frames that were taken at exactly 60-second intervals.

### Cell behavior scoring

Each frame of each track was scored as polarized or unpolarized in a random order by a researcher blind to the experimental condition. Scoring was performed using the membrane slice taken close to the coverslip. Cells were considered to be unpolarized if they were ruptured, blebbing, symmetric on two or more axes, or had multiple lamellipodia or a perimeter with high-frequency curvature. A subset of frames was scored in triplicate, which demonstrated that the internal disagreement rate was 5%. All other frames were scored once. Size, speed, and shape measurements were only made on individual frames that scored as polarized.

### Cell shape analysis

Cell shape analysis was based on principal component analysis (PCA) of 2D cell outlines, as previously described (Pincus & Theriot, 2007). Images were processed into vector representation for PCA using custom code written in Matlab R2018b (available upon request, **Figure S3A,C**). All analyses were performed on stacks cropped to 300 x 300 pixels in the XY-plane, centered on the membrane channel centroid identified during tracking as described above. For XY-plane shape analysis, the z-stack in the cytoplasmic mCherry or membrane channel (CellMask Deep Red) was converted into a maximum-intensity Z-projection, gamma-corrected, and segmented using an Otsu threshold. For XZ-plane shape analysis, the cytoplasmic mCherry channel was deconvolved with Huygens software (SVI) using a classic maximum-likelihood estimation and an empirically measured point-spread function. The deconvolved stack was gamma-corrected and segmented with Otsu thresholding. All segmented images were manually inspected using an overlay with the raw signal, and frames with large defects were removed from further analysis. Velocity alignment was performed on both the XY-plane segmentation and the 3D segmented stack. Briefly, the image or stack was rotated in the XY-plane, so that the instantaneous velocity vector was aligned parallel to the positive X-axis (i.e., pointing right horizontally, **Figure S3C**). The cells were translationally aligned by their centroid in all three dimensions, so that the shape analysis was conducted in the cell frame of reference. After velocity alignment, the 3D segmented stack was converted into an XZ-plane cross-section by making a maximum-intensity Y-projection. This XZ-plane projection may overestimate lamellipodial thickness because zebrafish keratocytes characteristically form wrinkles due to periodic detachment of the thin actin sheet from the substrate (Lou et al., 2021).

The aligned, binarized, and projected cell shapes from both the XY- and XZ-planes were then converted into 300-point contours, which outlined the boundary of the cell with point 1 assigned to the center of the cell front (as determined by where the velocity vector crossed the contour). These contours were outputted as 600-point vectors (consisting of the X- and Y-coordinates of each of the 300 points). Contour-vectors from all frames that scored as polarized were imported into CellTool (Pincus & Theriot, 2007) for two separate PCA runs, one for the XY-plane and one for the XZ-plane. Before-confinement and during-confinement cell shapes were pooled for their PCA runs (**Figure 3C**) and separate PCA runs were conducted with images pooled from before-hypertonic, during-hypertonic, and isotonic treatments (**Figure S4C,D**).

*In vivo* cell shapes were segmented manually using the quick selection tool in Photoshop 22.1.1 from cell clusters in the skin of mosaic fish during 0-10 minutes post wounding (*Microinjection for mosaic expression*, *Tissue laceration*). Basal cells (keratocytes) were imaged either using LifeAct-mNeonGreen or cytoplasmic mNeonGreen and membrane-targeted mRuby3 (**Figure S3B**). Movies were converted into maximum-intensity Z-projections. Although transgene-expressing cells tended to cluster together clonally, entire cell shapes, including cryptic lamellipodia, were visible in the LifeAct channel because the lamellipodia were brighter than neighboring cell bodies. In larvae expressing the cytoplasmic and membrane markers, lamellipodia could not be visualized beneath neighboring cells, so only the leading cell in each cluster was analyzed. These segmented images were naturally aligned so that the direction of the wound was parallel to the isolated keratocytes’ direction of instantaneous velocity. Therefore, the segmentations were converted directly to contours using Matlab and projected into the before/during-confinement shape space using CellTool.

Phase contrast images were segmented using the Directional Gradient Vector Flow Procedure (Seroussi et al., 2012), implemented using Matlab R2017b. For Figure 2C, each image shown in Figure 2B was registered to the masked bead using a custom Matlab script, then displayed as overlaid contours using CellTool.

### Volume and surface area measurements

Volume and surface area were measured from 3D reconstructions of spinning disk confocal images. Deconvolved and segmented 3D stacks of the cytoplasmic mCherry channel were prepared as described above (see *Cell shape analysis*). We tessellated the segmentations into triangulated surface meshes using custom code written in Python 3.7.1, using the *vtk* module and tools from the Allen Institute for Cell Science Spherical Harmonics Parameterization package (Viana et al., 2021). The volume and surface area of the mesh are reported as the cell volume and surface area.

### Statistical details

Each larva or isolated cell sample was considered an independent biological replicate, and multiple cells were measured in each larva and sample as technical replicates. Individual frames were averaged to calculate cell measurements, and individual cells were averaged to calculate sample measurements. Then, measurements were plotted either on a per-frame (Figure 3D-E), per-cell (dots in Figure 4, Figure 5B-E), or per-sample basis (triangles in Figure 4-5), using a “SuperPlot” style (Lord et al., 2020).

For Figure 3D and Figure 5E, effect sizes along each shape mode were calculated from sample measurements, after eliminating unpaired cells and samples with <3 cells, and are shown in Tables S1 & S3. For Figure 3E Welch’s ANOVAs with Games-Howell post-hoc tests were calculated along both shape modes because of unequal sample variances. Tests were performed on sample/larva averages and the resulting p-values and effect sizes are given in Table S2. For Figure 4A-D and Figure 5B-D, a paired t-test was performed on sample measurements, which were calculated by eliminating unpaired cells and samples with <3 cells, then sample-averaging the remaining paired measurements. P-values for the Pearson correlation coefficients (r) shown in Figure 4E-G were calculated as two-sided t-tests against the null hypothesis that r = 0 (i.e., that the two variables are uncorrelated), using Prism 9 (GraphPad).

## Data Availability

The data that support the results of this study are available from the corresponding author upon reasonable request.

