## Supplementary material for "Confined keratocytes mimic *in vivo* migration and reveal volume-speed relationship": S1

### Supplemental Figures

#### Figure S1

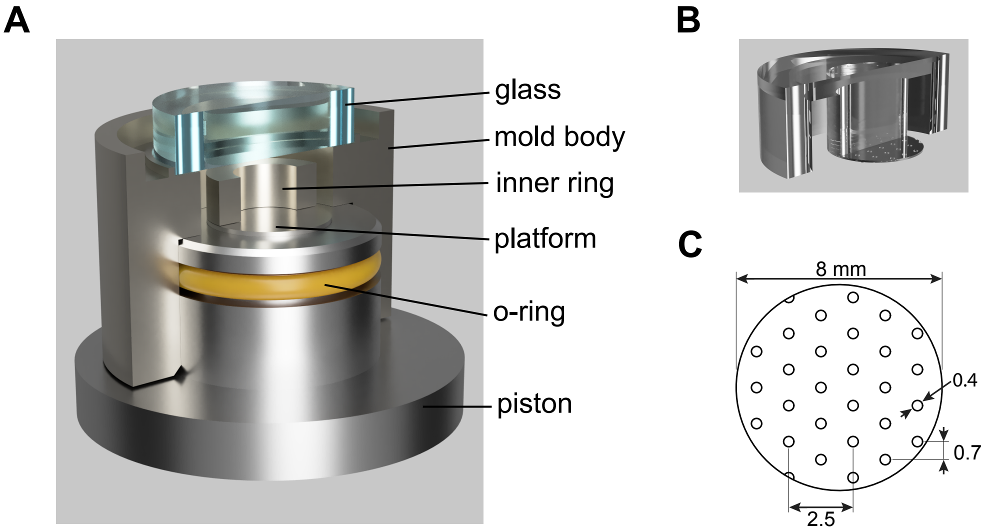

**Figure S1. Vacuum-controlled cell confiner design, based on** (Le Berre et al., 2014)**.**

**(A)** Rendering of the cell confiner mold showing intact piston, o-ring and platform. Mold body, inner ring and glass blank are cut in half for illustration purposes.

**(B)** Rendered PDMS cell confiner, cut in half, showing intact 8mm patterned glass coverslip. Overall size is 19 mm in diameter and 7 mm high.

**(C)** Glass coverslip showing pillar pattern. Pillars are 3.2 μm high. Dimensions in mm.

Figure S2

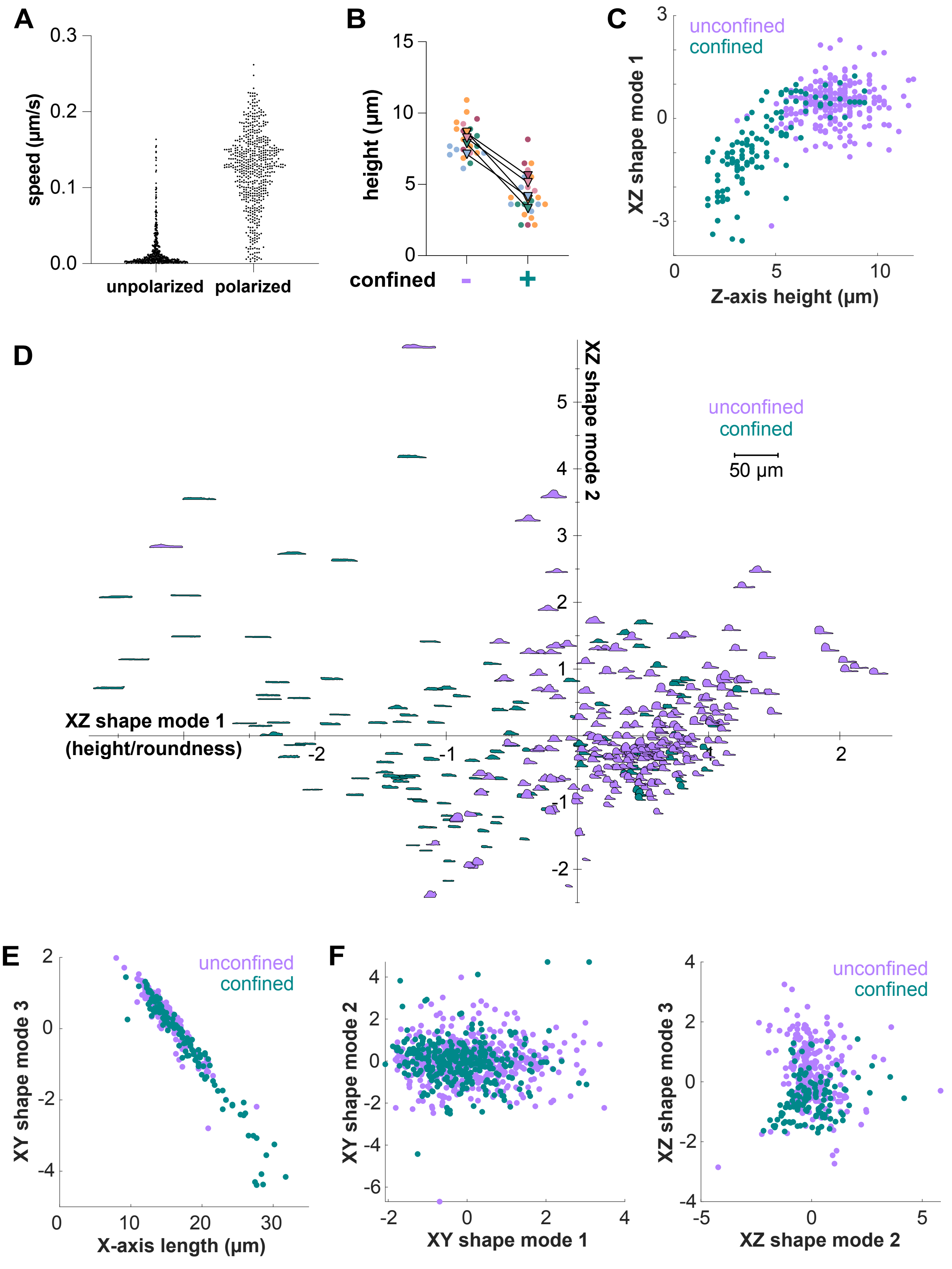
A
speed (µm/s)
0.3
0.2
0.1
0.0
unpolarized
polarized
B
15
10
5
0
height (µm)
confined
-
+
C
unconfined
confined
3
0
-3
XZ shape mode 1
0
5
10
Z-axis height (µm)
D
XZ shape mode 2
unconfined
confined
50 µm
XZ shape mode 1
(height/roundness)
E
unconfined
confined
XY shape mode 3
X-axis length (µm)
F
XY shape mode 2
XY shape mode 1
unconfined
confined
XZ shape mode 3
XZ shape mode 2

**Figure S2. Additional unconfined/confined behavior and shape results.**

**(A)** Plot of speed measurements grouped by polarization category. Data are pooled from unconfined and confined conditions. Each dot is an individual frame. Frames were randomly downsampled by half to assist with visualization ($n \geq 513$ cells per category shown).

**(B)** Plot of cell height grouped by confinement condition, as measured by the maximum height of a 3D segmented confocal stack. Dots indicate individual paired cell measurements from 5 samples. Sample averages indicated by triangles, with lines connecting the same sample across conditions. Cell and sample values are color-coordinated, with each color representing a different sample.

**(C)** Plot of cell height versus XZ shape mode 1 for all shapes in the XZ-plane shape space assembled from unconfined and confined conditions in the cell confiner experiments ($n > 120$ for each condition).

**(D)** Plot of all XZ cell contours in the XZ-plane shape space, positioned at their cell shape coefficients along XZ shape modes 1 and 2. The contours are scaled in both dimensions as indicated by the scale bar ($n > 120$ for each condition).

**(E)** Plot of cell length along the x-axis (the direction of migration) versus XY shape mode 3 for shapes across unconfined and confined conditions in the cell confiner experiments ($n > 120$ for each condition). Only mCherry-positive cells are shown to assist with visualization.

**(F)** Plots of top shape modes not included in Figure 3. Dots: observations of cell shape ($n > 120$ for each condition). Effect sizes for all shape modes are shown in Table S1.

Figure S3

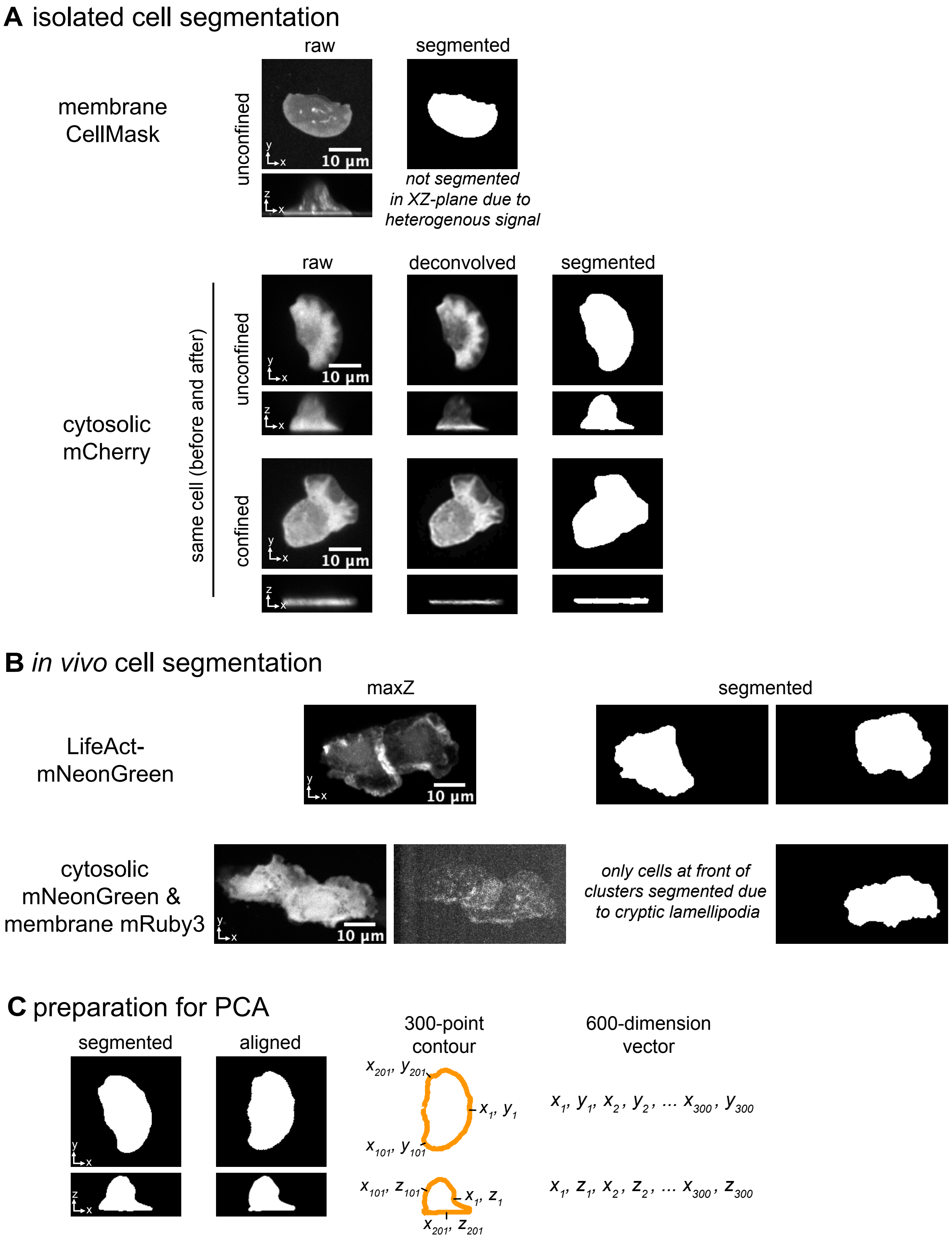
A isolated cell segmentation
raw
segmented
membrane CellMask
unconfined
10 µm
not segmented in XZ-plane due to heterogenous signal
raw
deconvolved
segmented
cytosolic mCherry
same cell (before and after)
unconfined
confined
10 µm
10 µm
B in vivo cell segmentation
maxZ
segmented
LifeAct-mNeonGreen
10 µm
cytosolic mNeonGreen & membrane mRuby3
10 µm
only cells at front of clusters segmented due to cryptic lamellipodia
C preparation for PCA
segmented
aligned
300-point contour
600-dimension vector
x201, y201
x1, y1
x101, y101
x1, y1, x2, y2, ... x300, y300
x101, z101
x1, z1
x201, z201
x1, z1, x2, z2, ... x300, z300

**Figure S3. Image processing for shape, volume and surface area measurements.**

**(A)** Segmentation workflow for isolated keratocytes. Samples with mCherry-negative cells were segmented in the XY plane using maximum-Z projections of CellMask Deep Red stacks. Samples with mCherry-positive cells were segmented in the XY-plane (using maximum-Z projections of mCherry stacks) or in 3D using deconvolved mCherry stacks. The 3D masks were projected into the XZ-plane for PCA-based shape analysis or converted to triangulated meshes for volume and surface area measurements. The confined mCherry cell is shown with a gamma correction for display purposes.

**(B)** Segmentation workflow for cells observed *in vivo* using larvae mosaically expressing either a LifeAct marker or cytosolic and membrane markers (*TgBAC(ΔNp63:Gal4)* larvae injected with *UAS:mNeonGreen-P2A-mRuby3-CAAX* or *UAS:LifeAct-mNeonGreen-P2A-mRuby3-CAAX* plasmid at the 1- or 2-cell stage).

**(C)** Vectorization workflow to prepare segmented masks for PCA-based shape analysis, shown using the unconfined mCherry cell shown in (A). Centroid-centered masks were rotated in the XY plane so that the instantaneous cell velocity pointed in the positive x-axis, then converted to 300-point contours with point 1 where the contour crossed the positive x-axis. Contour point coordinates were outputted in vector representation for analysis in CellTool.

Figure S4

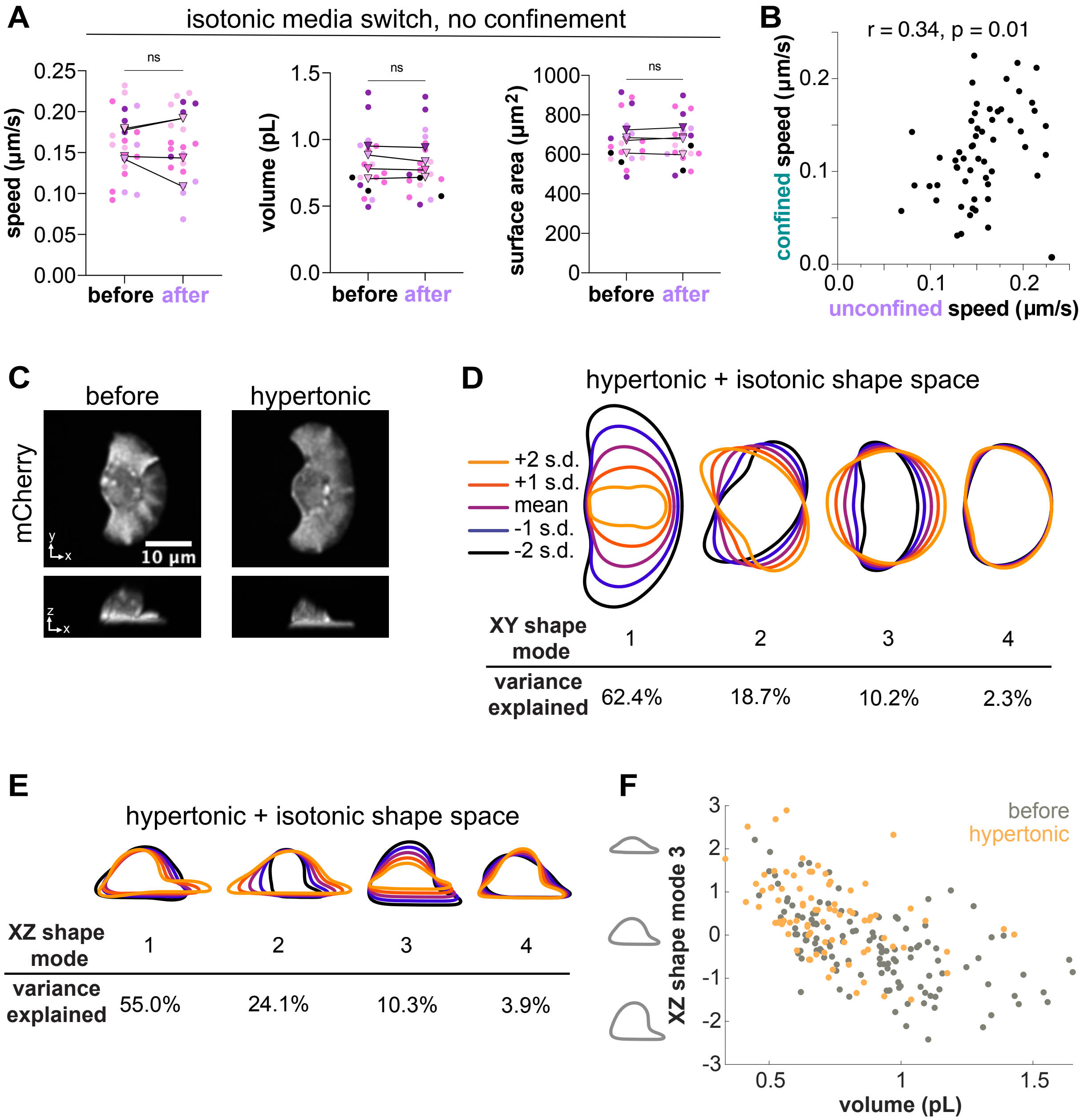

**Figure S4. Additional speed, size, and hypertonic shape analyses.**

**(A)** Control speed, volume, and surface area comparisons of cells before and after a media switch to identical, isotonic medium. Measurements made as described for the hypertonic shock experiment described in Figure 5B-D. Paired t-tests performed on 4 sample averages ($n \geq 3$ paired cells per sample), with $p > 0.05$ for all three measurements (ns: not significant).

**(B)** Plot of unconfined speed versus confined speed for cells in the PDMS cell confiner experiment. Each dot represents an individual cell ($n = 56$) from 10 samples. Pearson correlation coefficient (r) was calculated on a per-cell basis and the p-value is from a two-sided t-test against the null hypothesis that $r = 0$.

**(C)** Representative images of a keratocyte upon hypertonic shock. Keratocytes isolated from 2 dpf larvae expressing mCherry were imaged before (right image) and after (left image) the media was switched for media supplemented with 20-50 mg/ml sorbitol. Maximum-intensity projections of deconvolved spinning-disk confocal images.

**(D,E)** Modes of shape variation for the combined before-hypertonic, after-hypertonic, and control isotonic keratocyte population, as determined by principal component analysis (PCA) of velocity-aligned outlines. PCA performed on cells expressing mCherry, separately for 730 XY-plane outlines (D) and 674 XZ-plane outlines (E).

**(F)** Plot of cell volume versus XZ shape mode 3 for cells before and after hypertonic shock, showing a high correspondence between the two parameters.

Figure S5

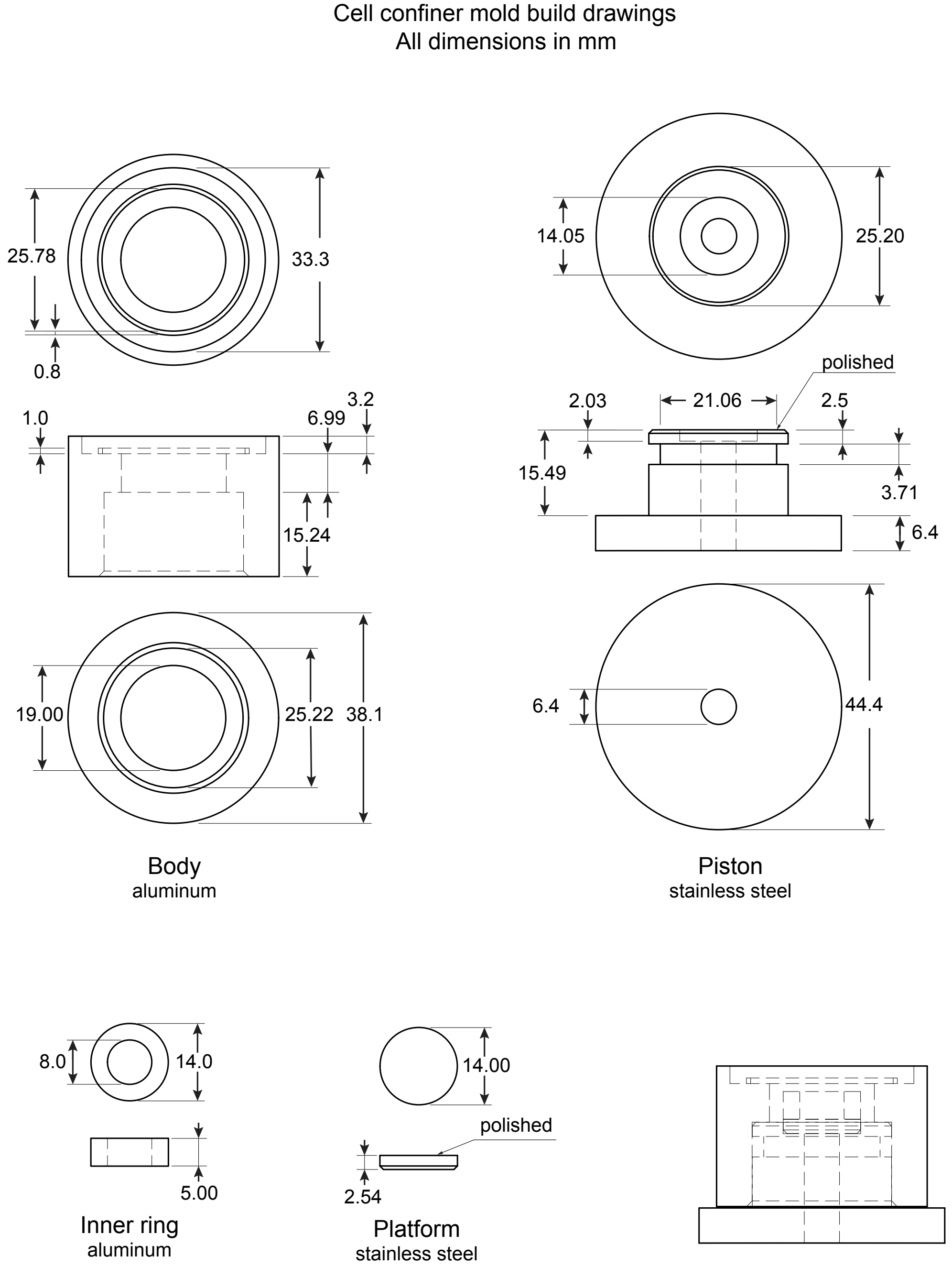

**Figure S5. Cell confiner mold build drawings.**

Exact drawings for machining the mold for the suction cup portion of the cell confiner. All dimensions are in mm. From top to bottom: body and piston top, cut-out, and bottom views; inner ring and platform top and cut-out views. Lower right: assembled mold cut-out view.

### Supplemental Tables

**Table S1. Effect size (Cohen's *d*) for unconfined versus confined shape mode coefficients.**

| Unconfined vs. Confined | | | | |
|---|---|---|---|---|
| XY plane | *Shape mode* | **1 (width)** | **2 (turning)** | **3 (inverse length)** |
| | *Cohen's d* | -0.41 | 0.12 | -2.0 |
| XZ plane | *Shape mode* | **1 (height)** | **2 (turning)** | **3 (size)** |
| | *Cohen's d* | -2.6 | 0.41 | -1.6 |

Cohen's *d* is the difference in means (confined minus unconfined) divided by the square root of the average sample variance. Calculated from 12 paired samples for the XY plane and 5 paired samples for the XZ plane, ignoring unpaired measurements on individual cells and samples with <3 cells with paired observations.

**Table S2. Intergroup p-values and Cohen's *d* for post-hoc Games-Howell statistical tests for Figure 3E.**

| XY shape mode 1 (width) | | | |
|---|---|---|---|
| | | **Confined (N=11)** | **In Vivo (N=4)** |
| **Unconfined (N=12)** | *p-value* | 0.18 | 0.0023 |
| | *Cohen's d* | -0.77 | -3.6 |
| **Confined (N=11)** | *p-value* | | 0.0064 |
| | *Cohen's d* | | -1.8 |
| XY shape mode 3 (inverse length) | | | |
| | | **Confined (N=11)** | **In Vivo (N=4)** |
| **Unconfined (N=12)** | *p-value* | 0.0020 | 0.011 |
| | *Cohen's d* | -2.7 | -5.8 |
| **Confined (N=11)** | *p-value* | | 0.015 |
| | *Cohen's d* | | -4.4 |

Tests performed on sample means for each dish or larva. Cohen's *d* is the difference in means (column minus row) divided by the square root of the average sample variance.

**Table S3. Effect size (Cohen's *d*) for unconfined versus confined shape mode coefficients.**

| Isotonic vs. Hypertonic | | | | |
|---|---|---|---|---|
| XY plane | *Shape mode* | **1 (width)** | **2 (turning)** | **3 (inverse length)** |
| | *Cohen's d* | -0.41 | -0.25 | -0.64 |
| XZ plane | *Shape mode* | **1 (height)** | **2 (turning)** | **3 (size)** |
| | *Cohen's d* | 0.62 | -0.11 | 1.5 |

Cohen's *d* is the difference in means (hypertonic minus isotonic) divided by the square root of the average sample variance. Calculated from 12 paired samples for the XY plane and 10 paired samples for the XZ plane, ignoring unpaired individual cells and samples with <3 cells with paired observations.

**Table S4. Primer sequences.**

| Primer | Description | Sequence (5'-3') |
|---|---|---|
| 1 | Q5 mutagenesis LifeAct, F | TCCAAGGAGGAGGGCGGCAGCGGCGGCGGC AGCGGCGGCATGGTGAGCAAGGGCGAG |
| 2 | Q5 mutagenesis LifeAct, R | GATGGACTCGAACTTCTTGATCAAGTCGGCCA CGCCCATCCAGCCTGCTTTTTTGTACAAAG |

### Video Legends

**Video S1. *In vivo* keratocytes migrate towards the wound using actin-rich lamellipodia.**

Cells migrating in the tailfin of a wounded 3 day post fertilization (dpf) larva expressing LifeAct-mNeonGreen mosaically in keratocytes. Laceration wound was to the right approximately 2 minutes earlier. Frames acquired every 30 seconds. Top: maximum-intensity Z-projection. Bottom: reslice cross-section taken at the position indicated by the white line. The cells lower in the Z axis are on the opposite side of the tailfin.

**Video S2. Isolated keratocytes migrate persistently using actin-rich lamellipodia.**

Isolated keratocyte expressing LifeAct-GFP and migrating randomly and persistently on a collagen-coated glass coverslip. Frames acquired every 10 seconds. Top: maximum-intensity Z-projection. Bottom: reslice cross-section taken at the position indicated by the white line. Images are gamma-corrected to better visualize the entire cell and binned by a factor of two using averaging for better comparison with Video S1.

**Video S3. Isolated keratocytes rapidly change shape in areas of reduced confinement.**

Isolated keratocytes were plated beneath an agarose gel overlay, along with rigid beads to hold up the agarose like tent-poles. This cell was initially observed far from a bead and displays rapid shape change as it approaches the area of reduced confinement around the bead. Phase contrast images taken every 5.5 seconds.

**Video S4. Isolated keratocytes rapidly change shape upon vacuum-controlled confinement.**

Isolated keratocytes labeled with CellMask Deep Red were plated in the vacuum-controlled PDMS cell confiner. This cell was initially observed with the ceiling in the raised position (>50 μm). As the video progressed, the vacuum was increased, lowering the ceiling to a height of 3 μm. Left: phase contrast images taken every 10 seconds. Right: CellMask stacks taken every 30 seconds; top: maximum-intensity Z-projection; bottom: maximum-intensity Y-projection, with a gamma correction to visualize the entire cell.
